# *S*-nitrosoglutathione reductase acts as a mitophagy receptor to control mitochondrial quality in plants

**DOI:** 10.64898/2026.10.28.758855

**Authors:** Huifang Lin, Hongyan Guo, Rong Wu, Wan Wang, Peng-Fei Jia, Shina Sun, Ru Bie, Hong-Ju Li, Wei-Cai Yang, Jianru Zuo

## Abstract

Mitochondria are central for cellular energy metabolism, and their quality is warranted by multiple mechanisms, including mitophagy, a process of selectively removing damaged, aged, or excess mitochondria. However, the regulatory mechanism of mitophagy remains elusive in plants. Here, we report that Arabidopsis *S*-nitrosoglutathione reductase 1 (GSNOR1), an enzyme that has long been known to degrade *S*-nitrosoglutathione, interacts with the outer mitochondrial membrane protein Voltage-Dependent Anion Channel 3 (VDAC3) to mediate mitophagy. GSNOR1 translocates to depolarized mitochondria, binds to VDAC3, and interacts with Autophagy-related 8 (ATG8) via its ATG8-interacting motifs, thereby engulfing damaged mitochondria in autophagosomes for mitophagic degradation. Mutations in *GSNOR1* or *VDAC3* cause the accumulation of misshapen mitochondria and hypersensitivity to mitochondrial stress. We propose that GSNOR1 acts as a cargo receptor interacting with VDAC3 and ATG8 to target depolarized mitochondria to mitophagy, thereby regulating cellular homeostasis.

## Main

Autophagy is a cellular process that targets macromolecules or organelles to the double-membraned autophagosome, followed by fusing to the lysosome or vacuole where the targets are degraded into small molecules for recycling. This highly conserved process is vital for both cellular homeostasis and survival under stress and starvation conditions. Autophagy is generally classified into three types based on the different pathways of substrate transport to lysosomes or vacuoles, namely macroautophagy, microautophagy, and molecular chaperone-mediated autophagy (Gross et al., 2025; Kuchitsu and Taguchi, 2024; Liu and Bassham, 2012; Marshall and Vierstra, 2018). Depending on the availability of nutrients, autophagy can also be classified into starvation-induced non-selective autophagy and selective autophagy. While non-selective autophagy is a key strategy for survival under starvation conditions, selective autophagy occurs under nutrient sufficient conditions and is generally initiated in response to cellular and environmental stresses to maintain cellular homeostasis (Gross et al., 2024; Marshall and Vierstra, 2018; Park et al., 2023; Wang et al., 2026). In plants, autophagy is involved in regulating various physiological processes, including growth, development, and both biotic and abiotic stress responses (Gross *et al*., 2025; Liu and Bassham, 2012; Marshall and Vierstra, 2018; Wang *et al*., 2026).

In selective autophagy, a target or cargo destined for degradation is specifically recognized by a receptor protein that in turn interacts with Autophagy-related 8 (ATG8), resulting in the engulfment of the target in autophagosome and subsequent autophagic degradation. The interaction of a cargo or a receptor with ATG8 is mediated by ATG8-Interacting Motif (AIM; also known as LC3-interacting region or LIR motif) located in a β-sheet (Noda et al., 2010; North et al., 2025). A typical AIM/LIR consists of a core motif of four amino acid residues [W/F/Y]XX[L/I/V], where X stands for any amino acid (Johansen and Lamark, 2020; Noda *et al*., 2010; North *et al*., 2025). In addition to AIM, the receptor-ATG8 interaction has also been shown to be mediated by the Ubiquitin-Interacting Motif (UIM), characterized as Ψ-ζ-X-A-Ψ-X-X-S (Ψ stands for hydrophobic amino acids; ζ stands for hydrophilic amino acids; and X stands for any amino acid) (Marshall et al., 2019; Stephani and Dagdas, 2020).

Mitochondria are central for cellular metabolism and energy supplies. When challenged by nutrient starvation or stresses, excessive, aged, or damaged mitochondria are subjected to autophagic degradation, a process known as mitophagy, which acts as a key strategy to maintain mitochondrial homeostasis (Broda et al., 2018; Pickles et al., 2018; Yang et al., 2024). Defects in mitophagy have been associated with abnormal growth and development as well as a variety of diseases in humans (Broda *et al*., 2018; D’Arcy, 2024; Pickles *et al*., 2018; Yang *et al*., 2024). In plants, mitophagy is largely induced by nutrient starvation, senescence, and mitochondrial stresses, and is involved in regulating growth, development, and stress responses (Broda *et al*., 2018; Duckney et al., 2024; Kacprzak and Van Aken, 2022). Mitophagy is initiated by recognizing and binding of damaged or depolarized mitochondria by autophagic cargo receptors, followed by the formation of autophagosome or mitophagosome. In principle, a mitophagy cargo receptor should contain an AIM motif, a UIM motif, or other types of motifs capable of binding to ATG8, and should localize at or associate with mitochondria. These features allow a cargo receptor recognizing damaged mitochondria. A number of mitophagy receptors have been identified in yeast and animals (Cen et al., 2020; Kanki et al., 2009; Liu et al., 2012; Okamoto et al., 2009; Sandoval et al., 2008; Wei et al., 2017; Yamashita et al., 2024). Recent studies have identified several putative receptors or receptor-like proteins for mitophagy in Arabidopsis (Li et al., 2022; Schmitt and Van Aken, 2023; Yang et al., 2023). Among those, TraB proteins, with dual mitochondrial and endoplasmic reticulum (ER) localization, interact with ATG8 to control mitophagy. The *trb1 trb2* double mutant accumulates an increased number of damaged mitochondria and shows altered stress responses (Li *et al*., 2022). Moreover, a mitochondrial E3 ligase SPL2 is found to degrade two mitochondrial outer membrane proteins that interact with an ER protein to mediate ER-mitochondrial interaction during seedling emergence (Tian et al., 2026). When challenged by carbon starvation, the mitochondria-localized FCS-like zinc finger (FLZ) proteins negatively regulate starvation-induced autophagy by inhibiting the activation of SnRK1, a key metabolic sensor (Yang *et al*., 2023). As FLZ proteins interact with ATG8 via the canonical AIM motifs, they may act as receptors for carbon starvation-induced mitophagy (Schmitt and Van Aken, 2023; Yang *et al*., 2023). Also in Arabidopsis, Friendly Mitochondria (Friendly or FMT), a homolog of Clustered Mitochondria Protein in animals, interacts with ATG8 and is possibly localized to damaged mitochondria during mitophagy (El Zawily et al., 2014; Kacprzak and Van Aken, 2023; Ma et al., 2021). FMT is proposed to act as a potential cargo receptor or to mediate mitophagy of depolarized mitochondria by preventing mitochondrial clustering to render autophagosomes more effectively capturing damaged mitochondria (Kacprzak and Van Aken, 2023; Ma *et al*., 2021; Schmitt and Van Aken, 2023). During later stage of mitophagy, mitochondrial fission and subsequent formation of mitophagosome require coordinated action of several key players, including ELM1, DRP3, and SH3P2 in response to heat stress (Ma et al., 2025a). Notably, core regulatory components of autophagy, including those for initiation, elongation and translocation, are highly conserved in eukaryotes. However, cargo receptors specifically targeting damaged mitochondria appear diverse in yeast, animals, and plants (Broda *et al*., 2018; Duckney *et al*., 2024; Schmitt and Van Aken, 2023).

Among numerous AIM-containing proteins, *S-*nitrosoglutathione reductase (GSNOR) is of particular interest. GSNOR proteins are highly conserved from bacteria to humans that irreversibly degrade *S-*nitrosoglutathione (GSNO), a major bioactive and reservoir form of nitric oxide (NO) (Liu et al., 2001; Sakamoto et al., 2002). Thus, GSNOR plays a vital role in regulating intracellular GSNO levels, thereby controlling downstream signaling processes mediated by NO and protein *S-*nitrosylation as a denitrosylase (Feng et al., 2019; Li et al., 2021). Loss*-*of-function mutations in Arabidopsis *GSNOR1*, known as *gsnor1*/*sensitive to hot temperatures5* (*hot5*)/*paraquat resistant2* (*par2*), cause severe growth and developmental defects (Chen et al., 2009; Feechan et al., 2005; Kwon et al., 2012; Lee et al., 2008). As a key regulator of intracellular GSNO levels, the biochemical activity and stability of Arabidopsis GSNOR1 are in turn controlled by NO-mediated *S-*nitrosylation (Guerra et al., 2016; Zhan et al., 2018). In Arabidopsis, *S-*nitrosylation of GSNOR1 at a highly conserved cysteine promotes its selective autophagic degradation mediated by a highly conserved AIM motif recognized by ATG8 (Zhan *et al*., 2018). In mouse, GSNOR is suggested to regulate mitophagy as a denitrosylase by modulating the *S-*nitrosylation levels of key components of mitophagy (Cirotti et al., 2021; Rizza et al., 2018; Tang et al., 2023). Because of the presence of a functional AIM motif (Zhan *et al*., 2018), GSNOR1 may also act as a cargo receptor for selective autophagy of other yet unidentified clients.

In this study, we present evidence showing that Arabidopsis GSNOR1 acts as a cargo receptor binding to the outer mitochondrial membrane protein Voltage-Dependent Anion Channel 3 (VDAC3) and ATG8 to target depolarized mitochondria to mitophagy in response to mitochondrial stress.

## Results

### *GSNOR1* controls mitochondrial number and quality

Previous studies show that Arabidopsis GSNOR1 protein is degraded via selective autophagy (Zhan *et al*., 2018) and that mouse GSNOR protein is involved in regulation of mitophagy by modulating de*-*nitrosylation of key components associated with the autophagic degradation of mitochondria (Rizza *et al*., 2018). Because GSNOR1 contains a highly conserved AIM motif, which is critical for its own selective autophagy and also is a key feature of cargo receptors for selective autophagy, we reasoned that GSNOR1 might regulate mitophagy. To test this possibility, we first examined the mitochondrial phenotype of *gsnor1*. An analysis of the number of mitochondria, judged by the copy numbers of the mitochondrial genome-specific genes, revealed that the copy numbers of the tested mitochondrial genome-specific genes (*COX1*, *ATP1*, *RPS4*, and *NAD6*) in *gsnor1* were significantly higher than those in wild type (Fig. 1a). A marginally increased copy number of these genes was observed in *atg2* and *atg5*, two autophagy deficient mutants (Fig. 1a), consistent with the observation that those two mutants had more mitochondria than wild type in dark-grown cotyledons by a different assay (Ma *et al*., 2021). Moreover, the accumulation of VDAC (an outer mitochondrial membrane marker), COXII (an inner mitochondrial membrane marker), and mtHSP70 (a matrix mitochondrial marker) proteins was at a substantially higher level in *gsnor1* than that in wild type (Fig. 1b,c).

**Fig. 1:**
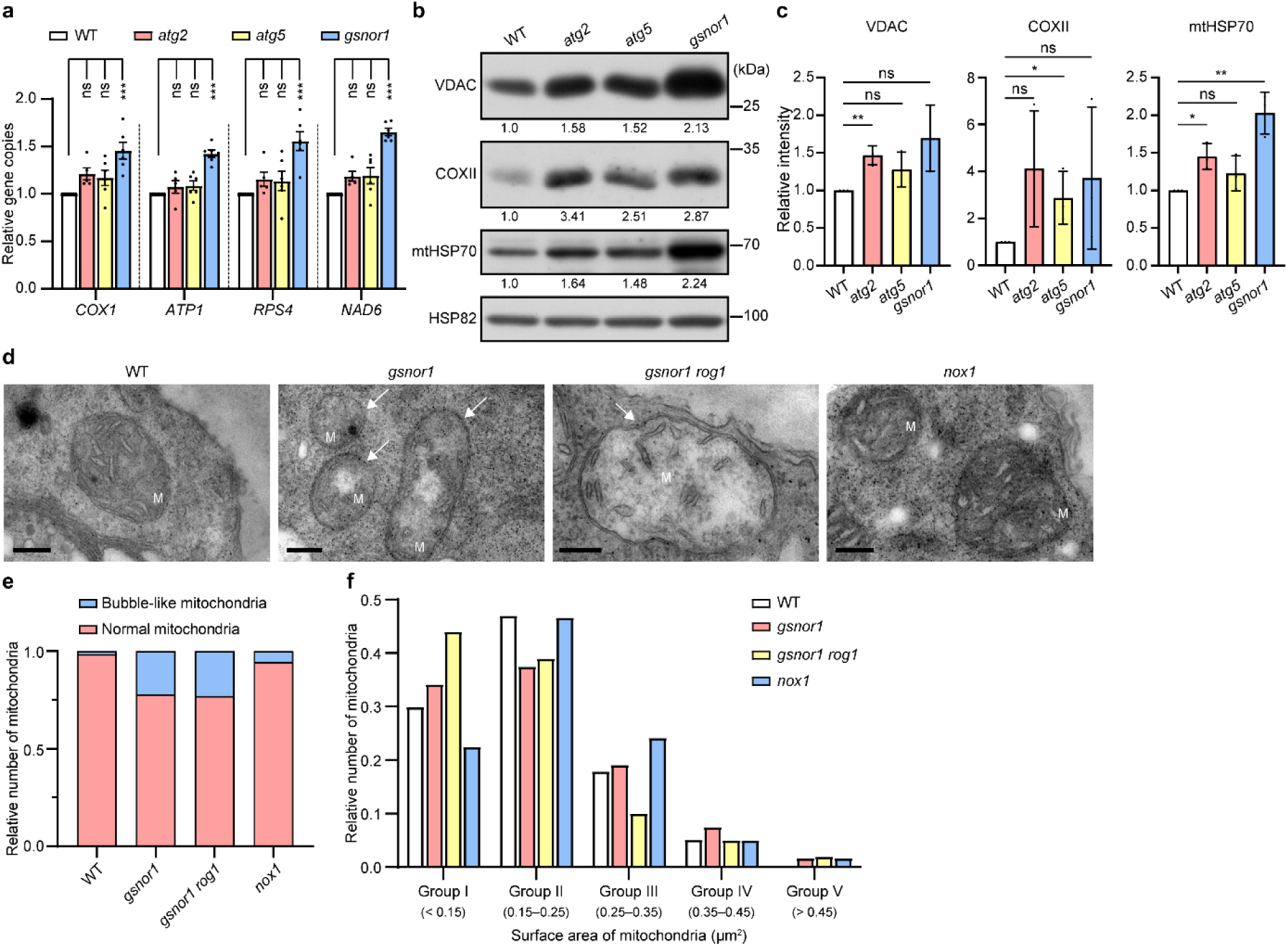
*GSNOR1* controls mitochondrial quality. **a**, Analysis of mitochondrial gene copy numbers of wild type (WT) and the indicated mutants by quantitative PCR (qPCR). DNA samples were prepared from 20-day-old seedlings. Bars represent SEM of more than five biological replicates. The nuclear-encoded single-copy gene *AtRpoTp* was used as an internal standard. Statistical significance was determined by one-way ANOVA followed by Tukey’s multiple comparisons test. ns, not significant; \*\*\**P* < 0.001. **b,** Immunoblotting analysis of mitochondrial marker proteins in 20-day-old seedlings with the indicated genotypes. HSP82 served as a loading control. Quantification is shown below the blot. The relative protein level in wild type is set as 1.0. VDAC: an outer mitochondrial membrane marker; COXII: an inner mitochondrial membrane marker; mtHSP70: a matrix mitochondrial marker. **c,** Quantitative analysis of the relative protein levels of VDAC, COXII, and mtHSP70 shown in (**b**). Data are presented as mean ± SD from three biological replicates. Statistical significance was determined by two-tailed Student *t*-test. ns, not significant; \**P* < 0.05; \*\**P* < 0.01. **d,** Transmission electron microscopy (TEM) of ultra-thin sections prepared from root tips of the indicated genotypes. White arrows denote damaged mitochondria not engulfed by autophagosomes. M, mitochondrion. Scale bars, 200 nm. **e,** Quantitative analysis of mitochondrial cristae morphology damage shown in (**d**) by counting the mitochondrial numbers of the defined phenotypes from multiple TEM micrographs. At least 100 mitochondria were counted for each sample. **f,** Relative numbers of mitochondrial surface area with the indicated ranges shown in (**d**). The analysis was performed by counting the mitochondrial numbers with the defined surface area ranges from multiple TEM micrographs. At least 100 mitochondria were counted for each sample.

To understand structural impacts of the *gsnor1* mutation, we examined the mitochondrial morphology by transmission electron microscopy of ultra-thin sections prepared from the root elongation zone. In wild type, spherical or rod-shaped mitochondria with well-organized cristae were observed. By contrast, irregular mitochondria with bubble-like cristae and the reduced number of cristae were found in *gsnor1* (Fig. 1d,e). Analysis of the surface areas of mitochondria showed that their sizes largely ranged smaller than 0.35 µm^2^ (Groups I-III), and a small population was larger than 0.35 µm^2^ (Groups IV and V). While the surface areas of mitochondria in wild type ranged between 0.15–0.25 µm^2^ (47.0%; Group II) and smaller than 0.15 µm^2^ (29.9%; Group I), the pattern was reversed in *gsnor1* with reduced population in Group II (37.5%) and increased population in Group I (34.2%) (Fig. 1f). The surface areas were marginally altered in Groups III and IV between wild type and *gsnor1* (Fig. 1f). Notably, Group V (larger than 0.45 µm^2^), constituted the smallest population, was nearly undetectable in wild type, but showed a significantly higher ratio in *gsnor1* (1.7%) (Fig. 1f), further suggesting that the *gsnor1* mutation causes irregular morphology. These results suggest that GSNOR1 is crucial for the maintenance of structure and membrane integrity of mitochondria.

Because GSNOR, a highly conserved enzyme catalyzing the degradation of GSNO, is an important regulator of NO homeostasis (Liu *et al*., 2001; Sakamoto *et al*., 2002), and Arabidopsis *gsnor1/hot5/par2* allelic mutants show a remarkably increased NO level (Chen *et al*., 2009; Feechan *et al*., 2005; Lee *et al*., 2008), we then asked if the observed abnormalities of mitochondria in *gsnor1* were associated with the increased NO level or alterations in protein *S*-nitrosylation levels. To address this question, we examined the mitochondrial phenotype of *nox1* and *gsnor1 rog1*, two mutants that both accumulate excessive amount of NO (Chen et al., 2020; He et al., 2004). *NO overproducer 1* (*nox1*) is characterized as a mutant containing an elevated NO level to repress floral transition (He *et al*., 2004). ROG1 (REPRESSOR OF GSNOR1) is proposed as a transnitrosylase catalyzing protein *S*-nitrosylation, and the *rog1* mutation restores defective growth and development of the *gsnor1* mutant to nearly normal, accompanying with proteome-wide reduction of protein *S*-nitrosylation in the *gsnor1* mutant background (Chen *et al*., 2020). Analysis of mitochondrial cristae morphology revealed that the *nox1* mutant largely showed a wild type-like phenotype with a higher ratio of the misshapen mitochondria (7/122; 5.7%) compared to wild type (2/136; 1.5%; Fig. 1d,e), suggesting that *NOX1* may play a minor role in mitochondrial quality control. By contrast, the *gsnor1 rog1* (23/100; 23.0%) double mutant displayed a phenotype similar to that of *gsnor1* (36/163; 22.1%; Fig. 1d,e). Consistent with these observations, *gsnor1 rog1* showed a phenotype of the mitochondrial area size similar to that of *gsnor1*, whereas *nox1* showed a pattern similar to that of wild type but opposite to that of *gsnor1* (Fig. 1f). These results suggest that NO and NO-mediated protein *S*-nitrosylation are not directly related to the dysfunctional mitochondria observed in *gsnor1*. Taken together, we conclude that *GSNOR1* plays an important role in regulating mitochondrial number and mitochondrial quality control.

### *GSNOR1* positively controls mitophagy

Data presented above show that the *gsnor1* mutant accumulates excessive number of damaged mitochondria, suggestive of impaired mitochondrial quality control. Under nitrogen-deficient conditions, the *gsnor1* mutant showed a phenotype similar to that of wild type, including significantly inhibited growth and non-specific degradation of inner and outer mitochondrial membrane marker proteins, whereas the macroautophagy-related *atg5* and *atg7* mutants were more sensitive to nitrogen deficiency (Extended Data Fig. 1a-e). Dark-induced carbon starvation has been reported to trigger mitophagy (Kacprzak and Van Aken, 2022). No apparent phenotypic difference was observed between wild type and the *gsnor1* mutant under the assay conditions (Extended Data Fig. 1a-d). These results suggest that *GSNOR1* does not have detectable effects in nitrogen- or dark-induced specific macroautophagy under the assay conditions.

To explore possible regulatory role of GSNOR1 in mitophagy, we tested the response of the *gsnor1* mutant to 2,4-dinitrophenol (DNP), a specific uncoupler of oxidative phosphorylation, which disrupts the electrochemical potential of the inner mitochondrial membrane, leading to the depolarized mitochondria targeting to degradation via *ATG5*-dependent mitophagy (Kacprzak and Van Aken, 2023; Li *et al*., 2022; Ma *et al*., 2021). Consistent with previous observations (Kacprzak and Van Aken, 2023; Li *et al*., 2022; Ma *et al*., 2021), when treated with DNP, the accumulation of the inner and outer mitochondrial membrane marker proteins (VDAC and COXII) was reduced in wild type, characteristics of the uncoupler-induced mitophagy. In the *atg5* and *gsnor1* mutants, while the accumulation of VDAC and COXII was comparable or slightly increased compared to that in wild type, the DNP-induced reduction of these proteins was compromised or slightly increased (Fig. 2a,b), suggesting that DNP-induced mitophagy is genetically dependent on *GSNOR1*.

**Fig. 2:**
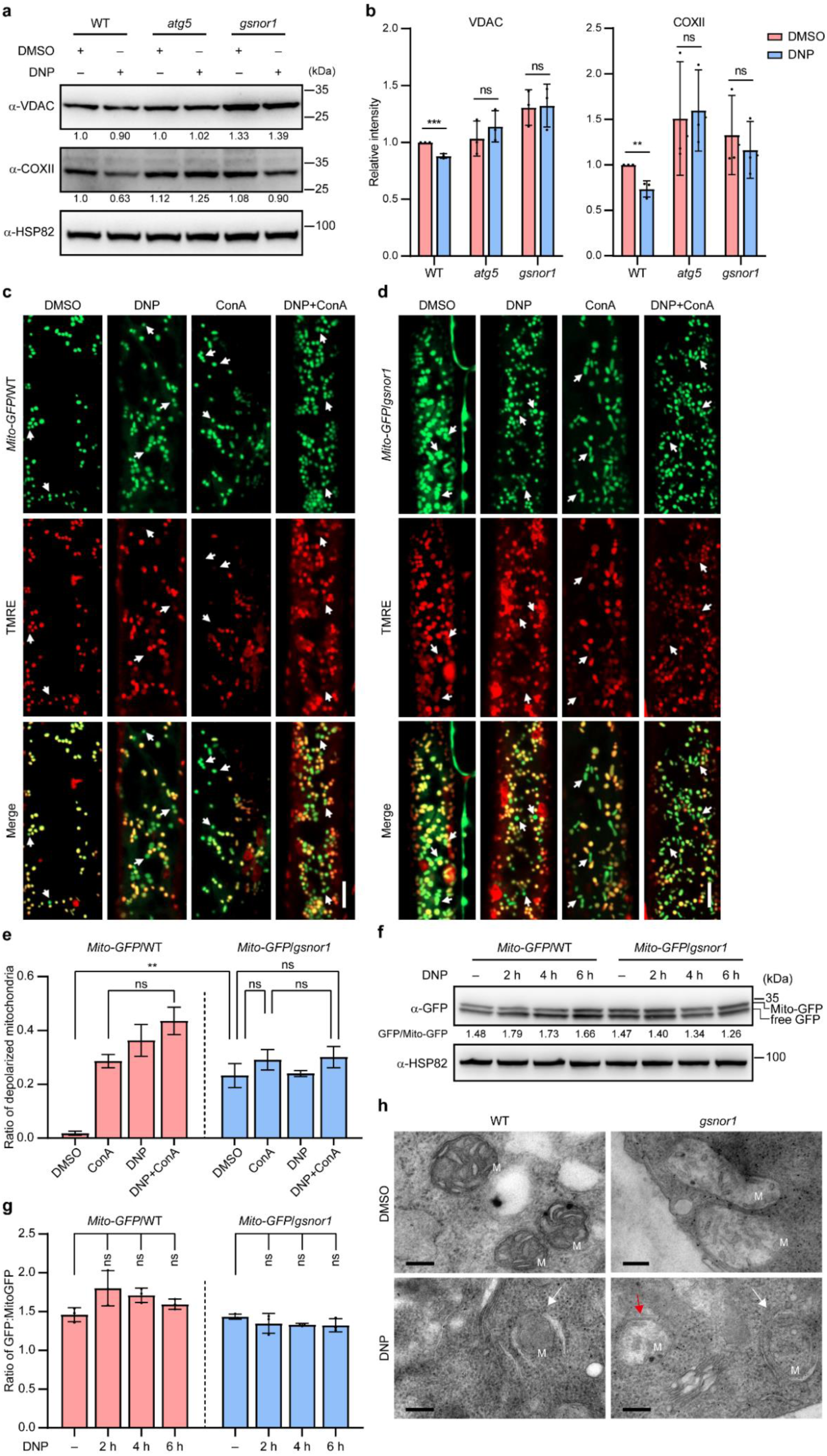
GSNOR1 positively regulates mitophagy. **a,** Immunoblotting analysis of mitochondrial outer membrane proteins (VDAC) and inner membrane protein (COXII) in 7-day-old seedlings of wild type (WT) and mutants with the indicated genotypes treated with DMSO (control) or 50 µM DNP for 4 h. HSP82 served as a loading control. **b,** Quantitative analysis of the relative protein levels of VDAC and COXII shown in (**a**). Data are presented as mean ± SD from three biological replicates. Statistical significance was determined by two-tailed Student *t*-test. ns, not significant; \*\**P* < 0.01; \*\*\**P* < 0.001. **c,d,** Confocal microscopy of mitochondria in roots derived from 7-day-old transgenic seedlings carrying a mitochondrial marker gene *Mito-GFP* in the wild type (**c**) and *gsnor1* (**d**) genetic backgrounds. The samples were treated with DMSO (control; 4 h), ConA (1 µM, 4 h), DNP (50 µM, 1.5 h), or ConA + DNP (4 h), followed by staining with TMRE (0.5 µM, 10 min). White arrows denote representative depolarized mitochondria. Scale bars, 5 µm. **e,** Quantitative analysis of depolarized mitochondria shown in (**c**) and (**d**). At least 1,000 mitochondria from 8 cells (three root samples) were analyzed. Data are presented as mean ± SEM. Statistical significance was determined by one-way ANOVA followed by Tukey’s multiple comparisons test. ns, not significant; \*\**P* < 0.01. **f,** Analysis of mitophagic flux in *Mito-GFP* transgenic seedlings in the wild-type (WT) and *gsnor1* genetic backgrounds. Cleavage of GFP was detected using an anti-GFP antibody. The upper and lower bands show full-length Mito-GFP and free GFP, respectively. Seven-day-old transgenic seedlings were treated with DMSO (control, 2 h) or with 50 µM DNP for the indicated times. Quantification was performed by calculating the ratio of free GFP to full-length Mito-GFP in each lane. HSP82 served as a loading control. **g,** Quantitative analysis of the ratio of free GFP to full-length Mito-GFP shown in (**f**). Data are presented as mean ± SD from three biological replicates. Statistical significance was determined by one-way ANOVA followed by Tukey’s multiple comparisons test. ns, not significant. **h,** Transmission electron microscopy of ultra-thin sections prepared from root tips derived from 7-day-old wild-type (WT) and *gsnor1* seedlings treated with 50 µM DNP for 4 h. White and red arrows denote damaged mitochondria engulfed and not engulfed by autophagosomes, respectively. M, mitochondrion. Scale bars, 200 nm.

To monitor dynamic changes of mitophagy in planta, we generated transgenic plants carrying a mitochondrial marker *Mito-GFP* (Kohler et al., 1997) in the wild type and *gsnor1* backgrounds, referred to as *Mito-GFP*/WT and *Mito-GFP*/*gsnor1*, respectively. A key hallmark of mitophagy is the depolarization of mitochondrial membrane, which is structurally distinguishable from normal mitochondria. Normal mitochondria, but not depolarized mitochondria, can be specifically labelled by tetramethylrhodamine ethyl ester (TMRE), a mitochondrial membrane potential-sensitive fluorescence dye (Qiao et al., 2025; Roussos et al., 2025; Zhou et al., 2016). Analysis of the root elongation zone showed that Mito-GFP and TMRE fluorescence signals were overlapped in nearly all foci in wild type (Fig. 2c), suggesting that the majority of mitochondria is normal. In the *gsnor1* mutant, a large portion of mitochondria was depolarized, showing largely as green foci when Mito-GFP and TMRE fluorescence signals were merged (Fig. 2d), consistent with the data presented above that the *gsnor1* mutation causes the accumulation of damaged mitochondria. When treated with DNP, depolarized mitochondria were dramatically increased in wild type; however, the *gsnor1* mutant, which contained a large portion of depolarized mitochondria, did not show apparent responses to DNP (Fig. 2c-e). These results suggest that the *gsnor1* mutation causes the accumulation of depolarized mitochondria in a manner similar to that induced by DNP. We noticed that *Mito-GFP*/WT plants had substantially less mitochondria than that observed in *Mito-GFP*/*gsnor1* plants (Fig. 2c-e), providing an additional line of evidence supporting a role of *GSNOR1* in controlling mitochondrial number.

To reinforce the above observations, the *Mito-GFP* transgenic plants were treated with concanamycin A (ConA), a vacuolar H^+^-ATPase inhibitor that blocks the autophagic flux, thus allowing the visualization of fluorescence in the vacuole (Tamura et al., 2003; Thompson et al., 2005). Treatment with ConA caused the accumulation of depolarized mitochondria in both wild type and *gsnor1* plants (Fig. 2c–e), suggesting that the vacuole is the final destination for the damaged mitochondria. Co-treatment of wild type with DNP and ConA showed similar phenotypes as treatments with individual reagents. However, the *Mito-GFP*/*gsnor1* plants had a high level of depolarized mitochondria, nearly comparable to that of DNP-treated wild type plants, and showed little or no response to DNP or ConA (Fig. 2c-e). In a mitophagic flux assay, the cleavage of Mito-GFP was enhanced upon DNP treatment in wild type, indicated by an increased GFP/Mito-GFP ratio; however, the cleavage was slightly suppressed in the *gsnor1* mutant, indicated by a deceased GFP/Mito-GFP ratio upon DNP treatment (Fig. 2f,g). Moreover, partial cleavage of GFP-TOM20-3 was detected in wild type, and the cleavage was enhanced by DNP treatment in wild type but not *gsnor1* (Extended Data Fig. 2a). These results suggest that the *gsnor1* mutation causes impaired mitophagy.

Transmission electron microscopy revealed that damaged mitochondria were engulfed in single-layer autophagosomes in wild type treated with DNP but not in the control samples (Fig. 2h). In the *gsnor1* mutant, while damaged mitochondria were found in both the control and DNP-treated samples, a significant portion of damaged mitochondria were not engulfed in autophagosomes under both control and DNP-treated conditions (Fig. 2h), suggesting that GSNOR1 plays a critical role in targeting damaged mitochondria to autophagosomes. Collectively, these results suggest that mitophagy is genetically dependent on *GSNOR1*.

### GSNOR1 interacts with ATG8 to engulf depolarized mitochondria to autophagosomes

Given the genetic dependency of mitophagy on *GSNOR1*, we next explored the function of GSNOR1 protein in the process. We generated transgenic plants carrying *pGSNOR1*::*GSNOR1-GFP* (referred to as *GSNOR1-GFP* hereafter) and *p35S*::*IDH1*-*mCherry* (a mitochondrial marker gene) transgenes in the *gsnor1* mutant background. Similar to the observation previously reported (Zhan *et al*., 2018), the *GSNOR1-GFP* transgene fully rescued the *gsnor1* mutant phenotype. Under normal growth conditions, GSNOR1-GFP distributed uniformly in the cytosol, barely co-localized with IDH1-mCherry (Fig. 3a). ConA treatment caused substantial co-localization of GSNOR1-GFP and IDH1-mCherry (Extended Data Fig. 2b). Similarly, when treated with ConA, co-localization of GSNOR1-GFP with MitoTracker Red-labeled mitochondria (Ma *et al*., 2021; Narendra et al., 2008) was readily detected (Extended Data Fig. 2c). When treated with DNP, GSNOR1-GFP was also observed in punctuated foci co-localized with IDH1-mCherry in a time-dependent manner (Fig. 3a,b), suggesting that GSNOR1 is translocated to depolarized mitochondria during the onset of DNP-induced mitophagy. When treated with both DNP and ConA to block the autophagic flux, the co-localization of GSNOR1-GFP and IDH1-mCherry formed enlarged focal structures that were readily detected (66.6% ± 14.8% co-localization; Fig. 3a). In a live imaging analysis, we found that GSNOR1-GFP was dynamically localized with the IDH1-mCherry-labeled mitochondria (Supplementary Video 1). Collectively, these results suggest that GSNOR1 is preferentially localized to depolarized mitochondria.

**Fig. 3:**
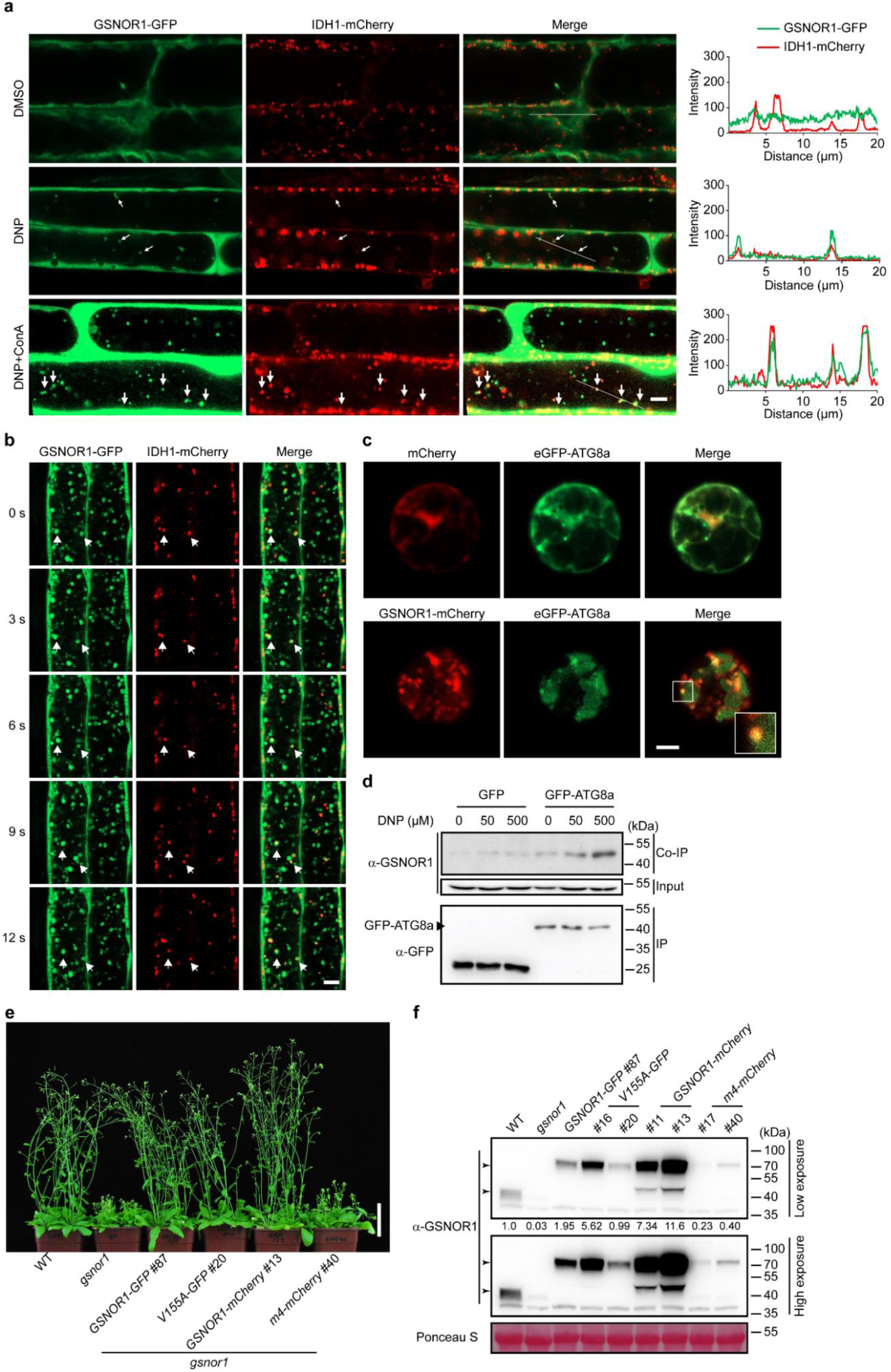
GSNOR1 localizes to depolarized mitochondria and interacts with ATG8a. **a,** Confocal microscopy of the root elongation zones derived from seedlings carrying both *GSNOR1-GFP* and *IDH1-mCherry* transgenes. Samples were treated with DMSO (control), DNP (50 µM), or DNP + ConA (1 µM) for 1.5 h prior to imaging. Line scans on the right side of the images show co-localization of GSNOR1-GFP (green) and IDH1-mCherry (red). White arrows indicate punctate co-localization signals. Scale bar, 5 µm. **b,** Confocal microscopy of the root elongation zones derived from 7-day-old seedlings carrying both *GSNOR1-GFP* and *IDH1-mCherry* transgenes. The samples were treated with ConA (1 µM) for 12 h prior to imaging at the indicated time points (second; s). White arrows indicate punctate co-localization signals. Scale bar, 5 µm. **c,** Confocal microscopy of Arabidopsis protoplast cells transformed with *GSNOR1-mCherry* and *GFP-ATG8a* genes. The boxed region shows an enlarged view at the bottom right shows co-localization of GSNOR1-mCherry and GFP-ATG8a. Scale bar, 5 µm. **d,** Analysis of the GSNOR1-ATG8a interaction by co-immunoprecipitation (Co-IP). Eight-day-old transgenic seedlings carrying *GFP* (control) or *GFP-ATG8a* transgenes were treated with the indicated concentrations of DNP for 2 h and then subjected to Co-IP assay. **e,** Six-week-old plants with the indicated genotypes grown under long-day condition (16-h light/8-h dark) in a growth chamber. Scale bar, 5 cm. **f,** Analysis of GSNOR1 protein in 6-week-old plants with the indicated genotypes by immunoblotting using an anti-GSNOR1 antibody. Equal loading was shown by Ponceau S staining. Quantification is shown below the blot. The relative protein level in wild type is set as 1.0. See also Extended Fig. 3d for the growth phenotype of other transgenic lines.

During autophagy, a key step is the link of ATG8 proteins to selective targets subjected to autophagic degradation. Previous studies showed that ATG8e interacted with GSNOR1 via an AIM (residues 152-155, YTVV) located in a β-sheet of GSNOR1 (Zhan *et al*., 2018). A closer inspection identified an additional highly conserved AIM-like motif in GSNOR1 (residues 269-272; FECI) located in a β-sheet (Extended Data Fig. 3a). A pull-down experiment showed that substitution mutations in both AIM motifs significantly reduced the interaction between GSNOR1 and ATG8e (Extended Data Fig. 3b,c). When transiently expressed in Arabidopsis protoplasts, co-localization of GSNOR1-mCherry and GFP-ATG8a, another member of the ATG8 family proteins, was observed (Fig. 3c and Extended Data Fig. 3c). In the *GFP-ATG8a* transgenic plants, DNP substantially enhanced the interaction between GSNOR1 and GFP-ATG8a in a dose-dependent manner as revealed by co-immunoprecipitation assay (Fig. 3d). These results suggest that GSNOR1 interacts with ATG8 proteins, targeting depolarized mitochondria to autophagosomes during mitochondrial stress.

To examine the physiological significance of these AIMs, we generated transgenic plants carrying double (*pGSNOR1*::*GSNOR1^Y152A^ ^V155A^ ^F269A^ ^I272A^-mCherry*; referred to as *GSNOR1^m4^-mCherry* hereafter) or single (*pGSNOR1*::*GSNOR1^V155A^-GFP*) mutations in the *gsnor1* background (we failed to obtained single mutants in the second AIM). While *GSNOR1^V155A^-GFP* fully rescued the developmental defects of *gsnor1* under the assay condition, *GSNOR1^m4^-mCherry* transgenic plants showed a *gsnor1*-like phenotype (Fig. 3e and Extended Data Fig. 3d), indicating that these two AIMs are essential for the GSNOR1 function. The nonfunctional GSNOR1^m4^-mCherry protein could be attributed to its instability (Fig. 3f). We noticed that the accumulation of GSNOR1^V155A^-GFP (line #20; relative level: 0.99) was comparable to that of endogenous GSNOR1 level (relative level: 1.0), approximately one-fold over that of GSNOR1^m4^-mCherry (line #40; relative level: 0.40) (Fig. 3f). However, these two transgenic proteins functioned completely different in rescuing the *gsnor1* mutant phenotype (Fig. 3e and Extended Data Fig. 3d), implying that the *GSNOR1^m4^* mutations in both AIM motifs may not only cause protein instability, but also cause the loss of the biochemical activity of the protein. The regulatory mechanism of GSNOR1^m4^ protein remains explored.

### GSNOR1 interacts with outer mitochondrial membrane-localized VDAC proteins

Proteins containing AIMs are considered as targets of selective autophagy and, in many cases, also act as receptors for cargos subjected to selective autophagy (Gross *et al*., 2025; Liu et al., 2024). Because GSNOR1 contains two functional AIMs and plays a critical role in mitophagy, we speculated that GSNOR1 might also act as a cargo receptor to mediate selective autophagy of mitochondria. If this is true, GSNOR1 should interact with a mitochondrion-localized protein, preferably localized at the outer membrane accessible by the cytosolic GSNOR1. In a yeast two-hybrid screen, we identified 25 putative GSNOR1-interacting proteins, including VDAC2 (1 clone) and VDAC3 (11 clones) (Supplementary Table 1). VDAC proteins, also known as porins, are a small family of highly conserved and abundant proteins localized on the outer mitochondrial membrane, which are also used as markers for the outer mitochondrial membrane (Doan et al., 2019; Kanwar et al., 2021; Najbauer et al., 2021; Oh et al., 2022; Shoshan-Barmatz et al., 2010) (Extended Data Fig. 4). In the Arabidopsis genome, 5 members have been identified, namely *VDAC1* through *VDAC5*, of which *VDAC5* was considered as a pseudogene (Lee et al., 2009; Tateda et al., 2011). The remaining four members of Arabidopsis VDACs showed 50%–68% homology. Notably, the positions of α-helices and β-sheets of VDACs are highly conserved across the plant kingdom and animal kingdom (Extended Data Fig. 4), implying that these structures are functionally important. Mutations in the four Arabidopsis *VDAC* genes (*VDAC1* through *VDAC4*) caused a variety of developmental abnormalities (Robert et al., 2012; Tateda *et al*., 2011).

When recapitulated in yeast cells, the interaction between GSNOR1/BD and VDAC3/AD was detected but not in other three VDAC proteins, and all VDAC/AD displayed varying degrees of auto-activated transcription activity (Extended Data Fig. 5a). In a pull-down experiment, the interaction of His-GSNOR1 recombinant protein with GST-VDAC recombinant proteins was detected at varying degrees, of which GST-VDAC3 showed the strongest interaction (Fig. 4a), consistent with the result that more VDAC3 clones were identified in the yeast two-hybrid screen (Supplementary Table 1). A co-immunoprecipitation experiment revealed that GSNOR1-FLAG was complexed with VDAC proteins as detected with polyclonal anti-VDAC antibodies that recognized VDAC1 through VDAC4 under normal growth conditions, and the interaction was enhanced by DNP (Fig. 4b). Moreover, the interaction of GSNOR1 with all four members of VDACs was also detected by the luciferase complementation experiment and bimolecular fluorescence complementation when transiently expressed in tobacco leaves (Fig. 4c,d). Notably, the co-localized GSNOR1 and VDAC proteins were observed in punctuated foci when transiently expressed in tobacco leaves and Arabidopsis protoplasts (Fig. 4d,e). These results suggest that GSNOR1 interacts with the outer mitochondrial membrane-localized VDAC family proteins, which may act to monitor the quality of mitochondria under normal growth conditions and promote mitophagy in response to stresses. Because VDAC3 showed the strongest interaction with GSNOR1, we analyzed VDAC3 in most experiments hereafter.

**Fig. 4:**
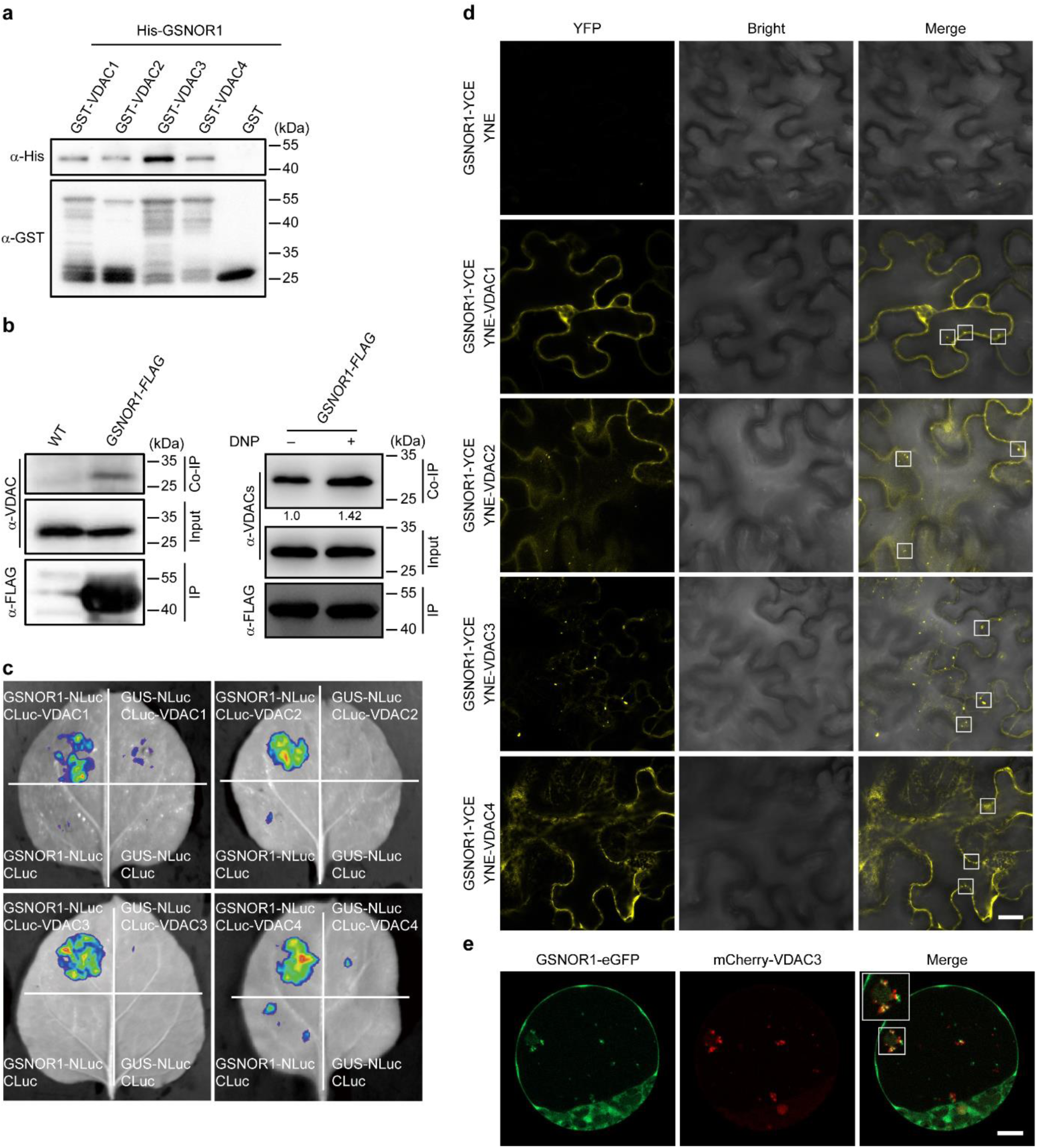
GSNOR1 interacts with VDAC proteins. **a,** Analysis of the interaction of His-GSNOR1 and GST-VDAC recombinant proteins by a GST pull-down assay. GST protein was used as a negative control. **b,** Analysis of the interaction between GSNOR1 and VDAC proteins by co-immunoprecipitation (Co-IP) in 8-day-old wild-type (WT) and *GSNOR1-FLAG* transgenic seedlings (left), and in *GSNOR1-FLAG* transgenic seedlings treated with or without 50 µM DNP for 2 h (right). Quantification is shown below the blot. The relative protein level in samples without DNP treatment was set as 1.0. The polyclonal anti-VDAC antibody recognizes VDAC1 through VDAC4 proteins. **c,** Analysis of the interaction of GSNOR1 and VDAC proteins by a luciferase complementation imaging assay in tobacco leaves infiltrated with *Agrobacterium* transformed with various combinations of transgenes as indicated. The assay was performed 2 days post-infiltration. **d,** Analysis of the interaction of GSNOR1 and VDAC proteins by a bimolecular fluorescence complementation assay in tobacco leaves infiltrated with *Agrobacterium* transformed with various combinations of transgenes as indicated. Confocal microscopy was performed 2 days post-infiltration. Co-localization signals are shown in boxed regions. Scale bar, 20 µm. **e,** Confocal microscopy of Arabidopsis protoplast cells transformed with *GSNOR1-GFP* and *FLAG-mCherry-VDAC3* transgenes. Co-localization signals are shown in enlarged views of the boxed regions. Scale bar, 10 µm.

### VDAC3 interacts with GSNOR1 and ATG8 via its N-terminal region

If VDAC proteins indeed play a role in mediating mitophagy, they should directly or indirectly interact with ATG8. In a pull-down experiment, we found that His-ATG8a recombinant protein directly interacted with GST-VDAC proteins (Fig. 5a). The interaction between VDAC and GFP-ATG8a proteins was also detected in a co-immunoprecipitation assay in the *GFP-ATG8a* transgenic plants (Fig. 5b). In a bimolecular fluorescence complementation assay, ATG8a was found to interact with VDAC proteins by forming punctuated co-localization signals (Fig. 5c and Extended Data Fig. 6a). When transiently expressed in Arabidopsis protoplasts, mCherry-VDAC3 and GFP-ATG8a were found to co-localize in punctuated foci (Fig. 5d). In a proteome-wide analysis, VDAC2 and VDAC3 were identified to interact with ATG8 via their UDS (UIM-docking site) and LDS (LIR/AIM-docking site) motifs, respectively (Marshall *et al*., 2019). Mutations in an LDS of ATG8a (Y65-L66) caused reduced interaction with both VDAC2 and VDAC3 (Extended Data Fig. 6b-d). However, mutations in a UDS of ATG8a (I92-F93-V94) showed enhanced interactions with VDAC2 and VDAC3 for unknown reasons (Extended Data Fig. 6b-d). These results suggest that VDAC3 proteins interact and co-localize with ATG8.

**Fig. 5:**
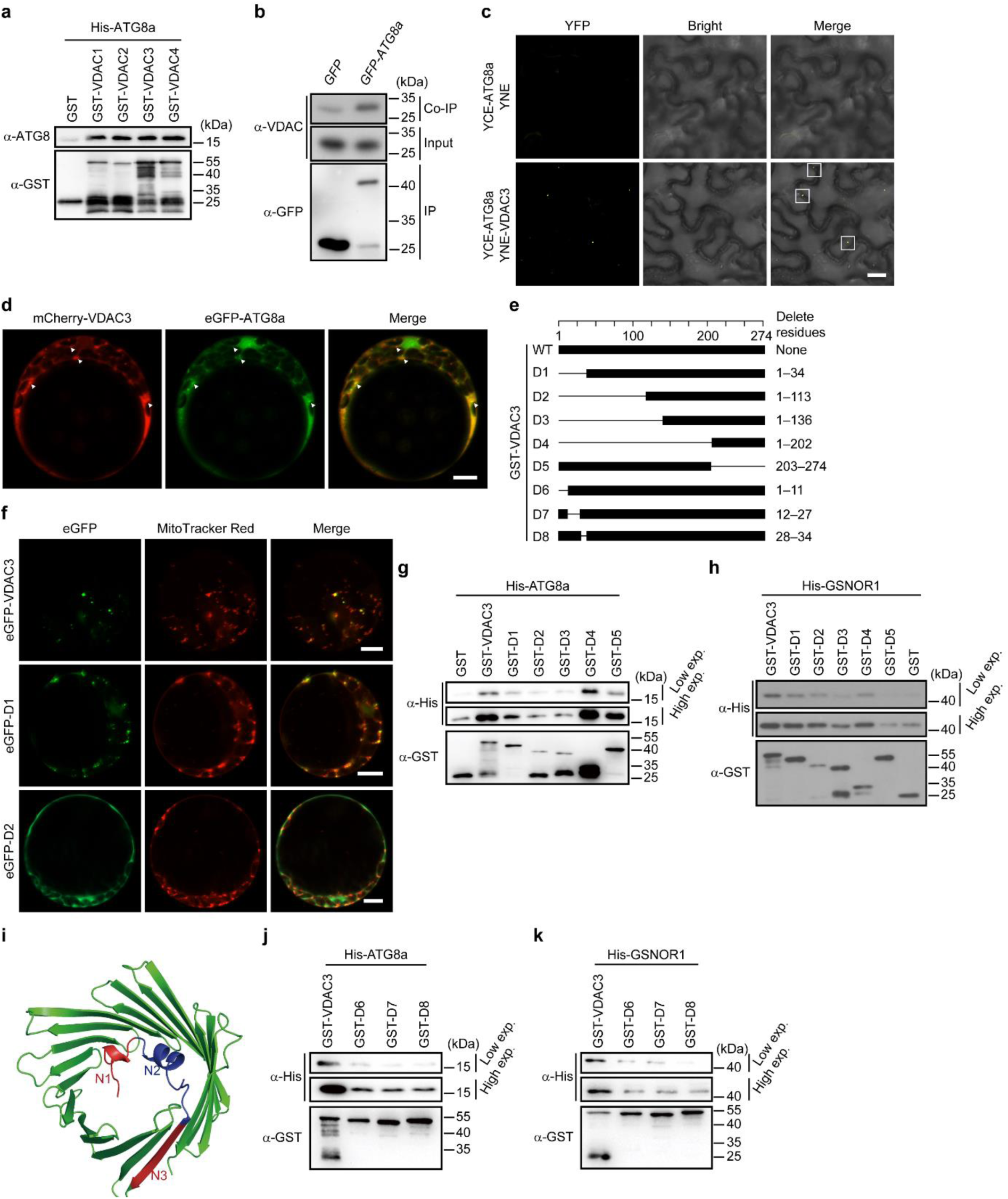
VDAC proteins interact with ATG8a and GSNOR1. **a,** Analysis of the interaction of GST-VDAC and His-ATG8a recombinant proteins by a GST pull-down assay. GST protein was used as a negative control. **b,** Analysis of the interaction of VDAC and ATG8a proteins by a co-immunoprecipitation (Co-IP) assay in 8-day-old seedlings carrying *GFP* or *GFP-ATG8a* transgenes. The polyclonal anti-VDAC antibody recognizes VDAC1 through VDAC4 proteins. **c,** Analysis of the interaction of VDAC3 and ATG8a proteins by a bimolecular fluorescence complementation assay in tobacco leaves infiltrated with *Agrobacterium* transformed with various combinations of transgenes as indicated. Confocal microscopy was performed 2 days post-infiltration. Co-localization signals are shown in the boxed regions. Scale bar, 20 µm. **d,** Confocal microscopy of Arabidopsis protoplasts transformed with *FLAG-mCherry-VDAC3* and *eGFP-ATG8a* transgenes. White arrowheads indicate punctate co-localization signals. Scale bar, 10 µm. **e,** Schematic representations of wild-type (WT) VDAC3 and deletion mutants fused to GST (GST not shown in the diagrams). **f,** Confocal microscopy of Arabidopsis protoplasts transiently expressing eGFP-VDAC3, eGFP-D1, or eGFP-D2. Protoplasts were stained with 200 nM MitoTracker Red for 15 min prior to imaging. Scale bars, 10 µm. **g,h,** Analysis of the interaction of wild-type (WT) GST-VDAC3 with D1 through D5 mutant recombinant proteins shown in (**e**) with His-ATG8a (**g**) or His-GSNOR1 (**h**) recombinant proteins by a GST pull-down assay. **i,** A projected 3-D structure of Arabidopsis VDAC3 protein trained with human hVDAC1 (PDB code: 2K4T) as a model. N1, N2, and N3 denote the putative α-helixes (N1 and N2) and a putative β-sheet (N3), of which are covered by deletional mutations D6-D8 shown in (**e**), respectively. **j,k,** Analysis of the interaction of wild-type (WT) GST-VDAC3 and D6-D8 mutant recombinant proteins shown in (**e**) with His-ATG8a (**j**) or His-GSNOR1 (**k**) recombinant proteins by a GST pull-down assay.

To map the regions interacting with ATG8a and GSNOR1, we generated truncated mutations of VDAC3 (Fig. 5e). When transiently expressed in Arabidopsis protoplasts, we noticed that VDAC-eGFP fusion proteins showed diffused subcellular localization patterns but not specifically localized in mitochondria for unknown reasons. We therefore used eGFP-VDAC fusion proteins in this study. We found that eGFP-VDAC3, eGFP-D1, and eGFP-D5 fusion proteins were localized to mitochondria in a punctate pattern and partially overlapped with MitoTracker Red, whereas eGFP-D2, eGFP-D3, and eGFP-D4 showed a diffuse cytosolic distribution (Fig. 5f and Extended Data Fig. 6e), suggesting that the D1 and D5 deletions do not have detectable effects on the subcellular localization of the fusion proteins. In a GST pull-down assay, the deletion of residues 1-34 (D1) dramatically reduced the interaction between His-ATG8a and GST-VDAC3 (Fig. 5g), suggesting that this region is important for the interaction. Consistent with this result, a recent study identified functional AIM motifs in the N-terminal region of VDAC3 (Ma et al., 2025b). Deletion of additional sequences (D2 and D3) showed a phenotype similar to that of D1. The D4 mutant, containing only the carboxyl terminal 41 residues (residues 203-274), showed the enhanced interaction with His-ATG8a, while the deletion of this region (D5) did not have detectable effect on the interaction (Fig. 5e,g). This result might be attributed to a cryptic signal or structure in D4. The interactions of His-GSNOR1 with these mutant proteins showed a pattern similar to that of His-ATG8a (Fig. 5h and Extended Data Fig. 5b). However, in contrast to that of His-ATG8a, D4 did not show an increased interaction with His-GSNOR1, implying that these two proteins may bind to VDAC3 with possibly different structural requirements. Based on the resolved 3-D structures of human and mouse VDAC proteins (Hiller et al., 2008; Ujwal et al., 2008), a putative structure model of Arabidopsis VDAC3 was constructed, in which the region spanning residues 1-34 contains two α-helices and the first β-sheet of a barrel structure that is formed with 19 anti-parallel β strands (Fig. 5i). To narrow down the interacting motifs of VDAC3, we made three additional mutants with smaller deletions, designated as D6 through D8, which covered the putative α1, α2, or β1 regions, respectively (Fig. 5e). All these three mutants showed substantially reduced interaction with His-ATG8a (Fig. 5j). Notably, these three mutants also nearly abolished their interactions with His-GSNOR1 (Fig. 5k). These results suggest that the amino terminal region (residues 1-34) of VDAC3 is critical for its interaction with both ATG8a and GSNOR1. Collectively, these results suggest that VDAC3 interacts with both GSNOR1 and ATG8a through the first 34 amino acid residues.

### The GSNOR1-VDAC3-ATG8 complex modulates mitochondrial stress responses

Data presented above showed that GSNOR1, ATG8a, and VDAC3 interact with one another. We next asked if these three proteins are in a same complex. To test this possibility, we co-transformed *FLAG-mCherry-VDAC3* and *HA-BFP-ATG8a* transgenes driven by a 35S promoter into protoplasts prepared from *GSNOR1-GFP* transgenic plants. Co-localization of these three proteins were detected in punctuated foci (Fig. 6a), indicating that these proteins form a complex in autophagosomes.

**Fig. 6:**
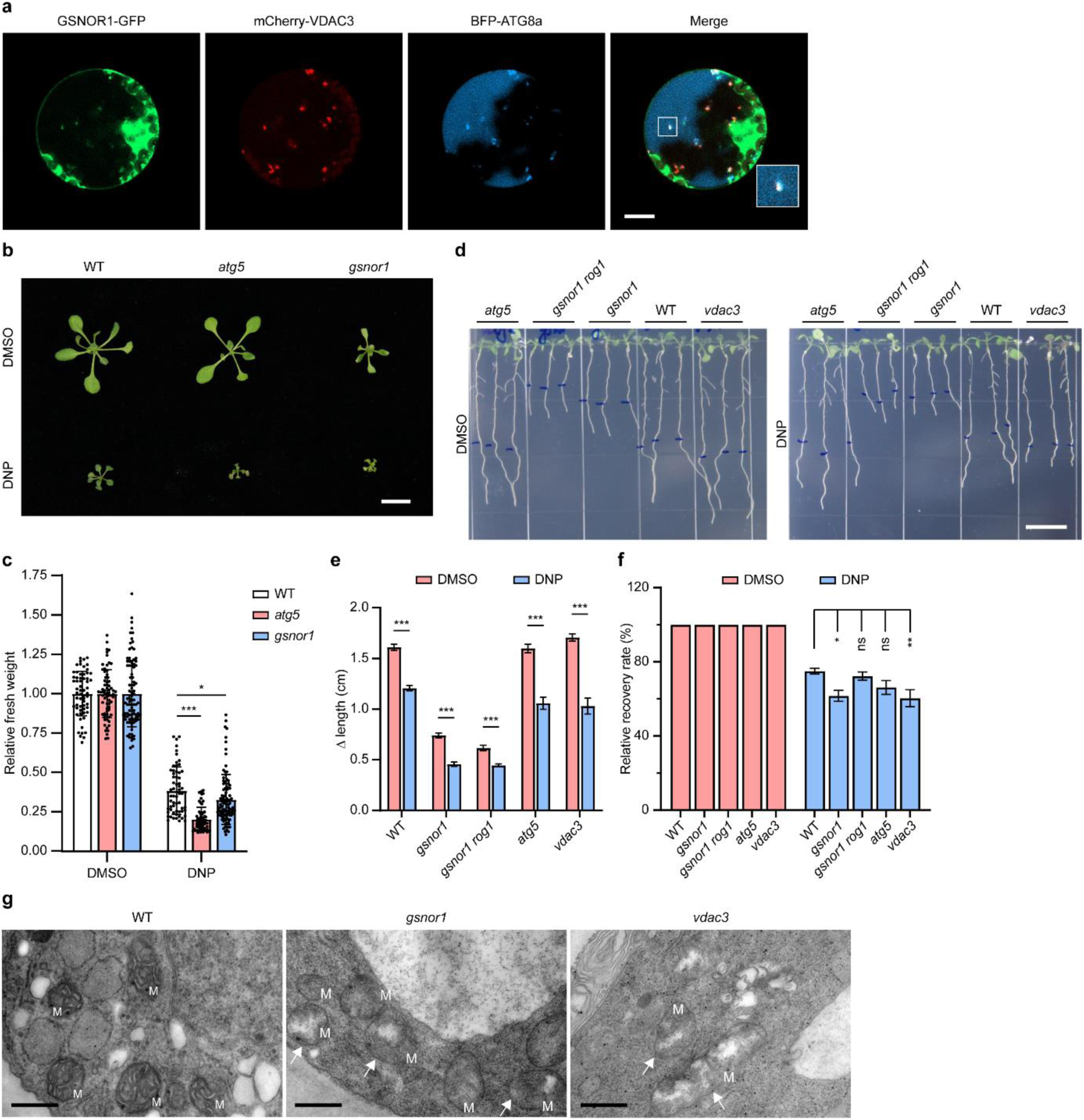
The GSNOR1-VDAC3-ATG8 complex modulates mitochondrial stress responses. **a,** Confocal imaging of Arabidopsis protoplasts prepared from *GSNOR1-GFP* transgenic plants transformed with *FLAG-mCherry-VDAC3* and *HA-BFP-ATG8a*. The boxed region shows an enlarged view of the co-localized signal. Scale bar, 10 µm. **b,** Seven-day-old wild type (WT), *atg5*, and *gsnor1* seedlings germinated and grown on 1/2 MS agar medium were transferred to 1/2 MS agar medium containing 10 µM DNP and cultured for additional 9 days. Scale bar, 1 cm. **c,** Quantification of fresh weight of seedlings shown in (**b**). The relative fresh weight of wild-type seedlings treated with DMSO was set as 1.0. Data are presented as mean ± SD (*n* ≥ 60). Statistical significance was determined by one-way ANOVA followed by Tukey’s multiple comparisons test. \**P* < 0.05; \*\*\**P* < 0.001. **d,** Seven-day-old wild-type and mutant seedlings with the indicated genotypes germinated and grown on 1/2 MS medium were transferred to liquid 1/2 MS medium containing 50 µM DNP and treated for 4 h. After the treatment, seedlings were transferred to 1/2 MS agar medium (the positions of root tips marked in blue) and cultured for additional 48 hr for recovery growth. Scale bar, 1 cm. **e,** Quantification of the root length of seedlings shown in (**d**). Data are presented as mean ± SEM (*n* ≥ 20). Statistical significance was determined by two-tailed Student *t*-test. \*\*\**P* < 0.001. **f,** Analysis of root recovery growth rate of seedlings shown in (**d**). The relative root recovery growth rate of seedlings cultured on medium containing DMSO (control) for each genotype is set as 100%. Root elongation from the marked root tip position was measured after 48 h of recovery growth. Data are presented as mean ± SEM (*n* ≥ 20). Statistical significance was determined by one-way ANOVA followed by Tukey’s multiple comparisons test. ns, not significant; \**P* < 0.05; \*\**P* < 0.01. **g,** Transmission electron microscopy of ultra-thin sections prepared from root tips derived from 9-day-old wild-type (WT), *gsnor1*, and *vdac3* seedlings. White arrowheads denote damaged mitochondrial with abnormal cristae. Scale bars, 500 nm.

The *gsnor1* mutant shows significantly altered responses to a variety of stress (Chen *et al*., 2009; Feechan *et al*., 2005; Lee *et al*., 2008). Given that GSNOR1 may act as a cargo receptor to mediate mitophagy, we examined the response of the *gsnor1* mutant to DNP. Seven-day-old seedlings grown under normal conditions were treated with 10 µM DNP for an additional 9 days, and the relative fresh weight was analyzed. Upon treatment with DNP, the relative fresh weight of the *gsnor1* and *atg5* mutants was significantly lower than wild type (Fig. 6b,c). In a growth recovery experiment, seedlings were treated with DNP for 4 hr and then transferred to normal growth conditions culturing for 48 hr. Analysis of root elongation post DNP treatment showed that *gsnor1* and *vdac3* mutants had lower recovery growth rate than that of wild type, while *gsnor1 rog1* and *atg5* mutants had marginally reduced recovery growth rates (Fig. 6d-f).

Consistent with the above results, transmission electron microscopy revealed severely damaged mitochondrial cristae in the *vdac3* mutant (Fig. 6g), similar to that observed in the *gsnor1* mutant. However, these damaged mitochondria were not frequently associated with double-membrane autophagic structures under the assay conditions. Taken together, these results suggest that the GSNOR1-VDAC3 complex regulates stress responses via mitophagy.

## Discussion

In this study, we present multiple lines of evidence, obtained from genetic, cell biological, and biochemical studies, showing that GSNOR1 may act as a cargo receptor docking damaged mitochondria for mitophagic degradation via interacting with VDAC3 and ATG8. First, mutations in *GSNOR1* and *VDAC3* cause a similar phenotype characterized as the accumulation of dysfunctional mitochondria and hypersensitive to the oxidative phosphorylation uncoupler DNP, thus demonstrating the genetic dependency of mitophagy on these two genes. Second, GSNOR1 dynamically and preferentially localizes to depolarized mitochondria, which is a prerequisite of a regulator for mitophagy, accompanying with the co-localization of GSNOR1, VDAC3, and ATG8a in autophagosomes. These observations directly link GSNOR1 and VDAC3 to ATG8-mediated mitophagy. Third, GSNOR1 interacts and co-localizes with both ATG8 and VDAC3 in autophagosomes, characteristics of a cargo receptor of selective autophagy. Based on these observations, we propose that GSNOR1 functions as a cargo receptor, interacting with the outer mitochondrial membrane protein VDAC3 and targeting the depolarized mitochondria to ATG8, eventually docking to autophagosomes for degradation (Fig. 7). These findings not only identify a previously unrecognized mechanism regulating mitophagy, a highly conserved process in eukaryotes, and also uncover an unexpected role of GSNOR1 that has long been known as an enzyme degrading GSNO.

**Fig. 7:**
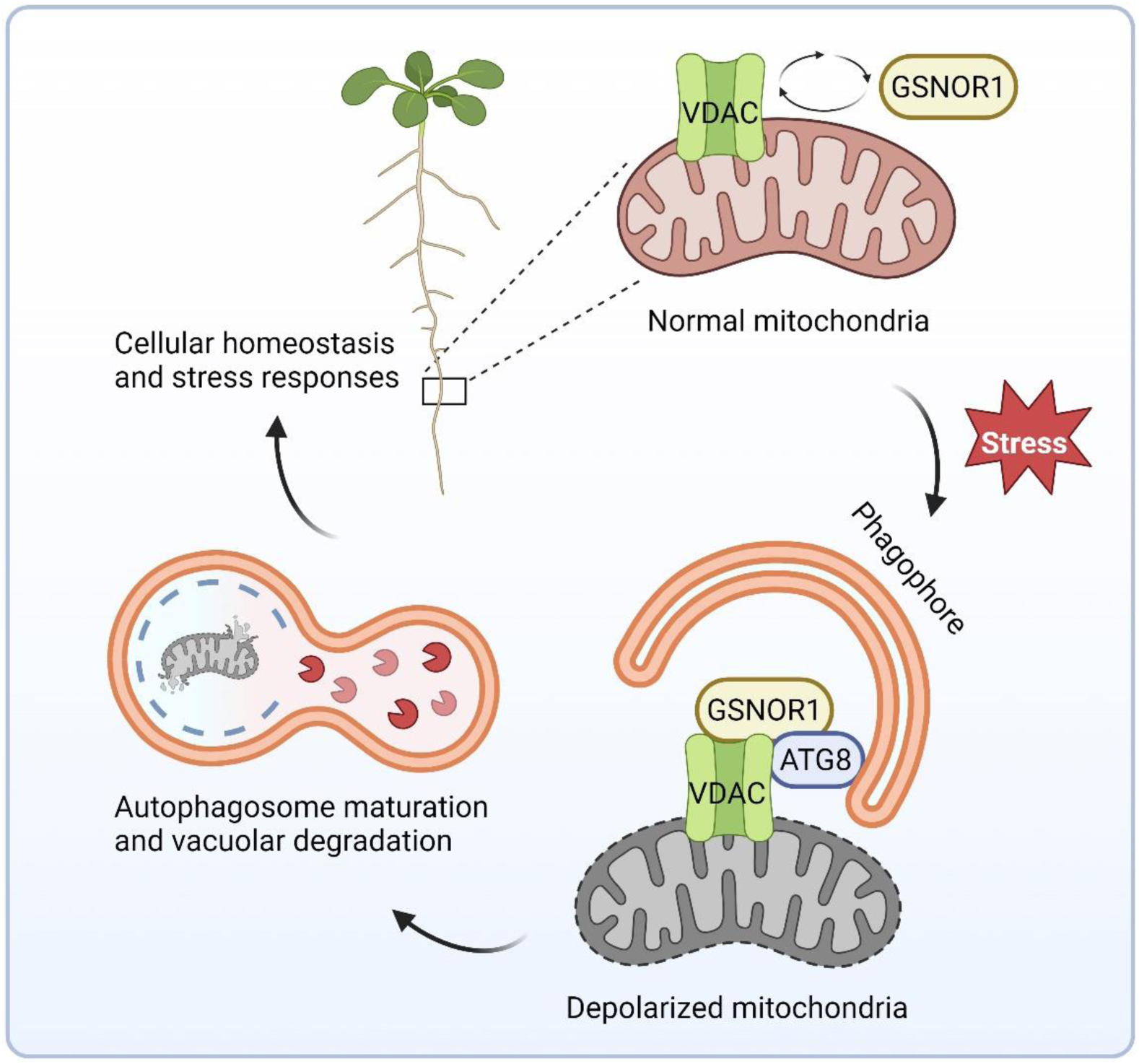
A model of GSNOR1-regulated plant mitophagy. Under normal growth conditions, GSNOR1 dynamically interacts with the outer mitochondrial membrane-localized VDAC3 protein, which may serve to monitor the quality of mitochondria. When challenged by stresses, mitochondria become depolarized, initiating the formation of a complex consisting of the mitophagy receptor GSNOR1, the phagophore-localized ATG8, and VDAC3. The GSNOR1-ATG8-VDAC3 complex recruits phagophore to encapsulate depolarized mitochondria, which subsequently fuses to vacuoles for autophagic degradation, thereby safeguarding cellular homeostasis and regulating stress responses.

VDACs are highly conserved proteins localized on the outer mitochondrial membrane. In animals, the leafhopper VDAC1 interacts with ATG8 to mediate mitophagy upon insect-borne virus infection (Chen et al., 2023) and the localization of the E3 ligase Parkin to depolarized mitochondria causes poly-ubiquitylation of VDAC1 at the onset of mitophagy (Geisler et al., 2010). Observations made in this study show that the Arabidopsis *vdac3* mutant is hypersensitive to DNP and has malformed mitochondrial structures, accompanying with the physical interaction of VADC3 with ATG8, implying a possible role of VADC3 in regulating mitophagy. Consistent with this notion, the Arabidopsis VDACs have been identified as ATG8-interacting proteins in a proteomic study (Marshall *et al*., 2019). Moreover, a recent study shows that VDAC1, VDAC2, and VDAC3 acts as mitophagy receptors recruiting ATG8 to depolarized mitochondria under DNP-induced mitochondrial stress (Ma *et al*., 2025b). Biochemically, VDAC3 directly interacts with both ATG8 and GSNOR1 via its amino-terminal region. The human VDAC1 (hVDAC) structure is characterized as a tunnel-like structure comprising 19 anti-parallel β-sheets, connecting to a short and flexible amino-terminal region (PDB code: 2K4T) (Hiller *et al*., 2008). Analog to the amino-terminal region of hVDAC1, a short stretch of Arabidopsis VDAC3 (residues 1-34) is critical for its interaction with both GSNOR1 and ATG8a, suggesting that GSNOR1 and ATG8a may bind to the outer mitochondrial membrane-localized VDAC3 in a cooperative fashion.

While it is not unexpected for the regulatory role of VDAC proteins (Broda *et al*., 2018; Geisler *et al*., 2010; Ma *et al*., 2025b), it is somewhat surprising to find that GSNOR1 functions as a receptor to mediate mitophagy, considering its canonical biochemical activity as an *S*-nitrosoglutathione reductase, also known as a glutathione-dependent formaldehyde dehydrogenase (Liu *et al*., 2001; Sakamoto *et al*., 2002). An interesting clue comes from the observation that while *HOT5/GSNOR1* is a key regulator of thermotolerance (Lee *et al*., 2008), heat stress induces piecemeal mitophagy (Ma *et al*., 2025a). However, it remains elusive if *HOT5/GSNOR1* is involved in piecemeal mitophagy. GSNOR1 contains two AIMs, one of which has been shown important for its own selective autophagy (Lubega et al., 2021; Zhan *et al*., 2018). Proteins containing functional AIMs are also considered as cargo receptors to target specific proteins or organelles to selective autophagy (Faruk et al., 2021; Stephani and Dagdas, 2020), and GSNOR1 is therefore a strong candidate as an autophagy receptor. While evidence presented in this study suggests that GSNOR1 acts as a cargo receptor to mediate mitophagy, it seems difficult to understand why a GSNO-degrading enzyme moonlights as a mitophagy receptor. Stress-induced burst of NO and reactive oxygen species (ROS) cause the damage of mitochondria (Dai et al., 2020; Huang et al., 2016; Zhang et al., 2022) and, on the other hand, *S*-nitrosylation (Guerra *et al*., 2016; Zhan *et al*., 2018) and oxidation (Sun et al., 2025) also negatively regulate the GSNOR1 activity. As a cytosol dominantly localized protein, GSNOR1 is presumably capable of rapidly sensing and subsequently responding to cellular redox alterations by adjusting its own enzymatic activity and evoking a mechanism clearing damaged cellular components in response to stresses. Echoed to this scenario, an analogous mechanism has been well appreciated in glyceraldehyde-3-phosphate dehydrogenase (GAPDH), a key enzyme catalyzing the production of 2,3-bisphosphoglyceric acid from glyceraldehyde-3-phosphate in glycolysis. In addition to its role in glycolysis, GAPDH is also involved in the regulation of a wide range of cellular activities, including autophagy, cell death, cell differentiation, and responses to carbon metabolic flux and other stimuli (Cornett et al., 2022; Dodson et al., 2013; Feng et al., 2024; Henry et al., 2015). Analogously, GSNOR1 may play dual and distinctive roles in NO signaling and mitophagy in a fashion similar to GAPDH. Moreover, considering the pleiotropic phenotype of the *gsnor1* mutant and the nature of a cargo receptor, it is predictable that GSNOR1 may function as a receptor of other yet unidentified clients during selective autophagy.

Given that GSNOR1 catalyzes the degradation of GSNO and that the *gsnor1* mutant accumulates excessive amount of NO and causes proteome-wide over-nitrosylation (Chen *et al*., 2020; Chen *et al*., 2009; Feechan *et al*., 2005; Hu et al., 2015; Lee *et al*., 2008), one would expect that NO or NO-mediated *S*-nitrosylation may also play an important role in regulating mitophagy. In this regard, a large portion of mitochondrial proteins have been identified as *S*-nitrosylated proteins by proteomics studies (Chang et al., 2014; Foster and Stamler, 2004; Hu *et al*., 2015), implying possible involvement of these potential targets in the NO-regulated mitophagy. A line of more direct evidence comes from the observation that the knockout of the mouse *GSNOR* gene, similar to that in Arabidopsis, also causes impaired mitophagy, which is attributed to excessive *S*-nitrosylation of Drp1 and Parkin (Rizza *et al*., 2018). Because both Drp1 and Parkin homologs are not found in plants, it is not likely a similar mechanism operating mitophagy in plant cells. Therefore, while the lack of the GSNOR activity evokes mitophagy in both mammals and plants, different mechanisms are found in these two systems, in which GSNOR acts as a denitrosylase and a cargo receptor, respectively. In fact, the observation that minimal mitochondrial phenotype is observed in the *nox1* mutant rules out the possibility of a major role of NO in regulating mitophagy under the assay conditions. Moreover, the *rog1* mutation genetically suppresses the *gsnor1* phenotype in multiple aspects, including the proteome-wide over-nitrosylation of *gsnor1*. However, the restoration or reduction of the *S*-nitrosylation level of *gsnor1* to near the wild-type level by *rog1* does not suppress the defective mitophagy phenotype, implying that neither NO nor NO-mediated protein *S*-nitrosylation plays a decisive role in regulating mitophagy under the assay condition. Despite of this notion, we do not exclude the possibility that GSNOR1 also functions as a denitrosylase to regulate mitophagy in plants. Instead, it will be of great interests to identify *S*-nitrosylated proteins regulating mitophagy in plants. Lastly, the interactions among ATG8, GSNOR1, and VADC3 are detected under normal growth conditions, supporting a notion that the machinery may act to monitor the quality of mitochondria under both permissive and nonpermissive growth conditions.

In conclusion, discoveries made in this study identify a unique and unexpected mechanism regulating mitophagy by GSNOR1 in plants. Because all major components in the pathway, including GSNOR, VDAC, and ATG8, are highly conserved, a similar regulatory scheme may also be functional to execute mitophagy in animals as well.

## Methods

### Plant materials and growth conditions

The Columbia-0 (Col-0) accession of *Arabidopsis* was used in this study. The *atg2*, *atg5*, *atg7*, *nox1*, and *vdac3* (SALK_127899) mutant seeds were provided by Dingzhong Tang, Yikun He, and ABRC, respectively (He *et al*., 2004; Wang et al., 2011). The *gsnor1* and *gsnor1 rog1* mutants have been previously described (Chen *et al*., 2020; Chen *et al*., 2009). Transgenic *Arabidopsis* plants were generated using the *Agrobacterium*-mediated floral dip transformation method (Bechtold and Pelletier, 1998).

Seeds were surface-sterilized and sown on 1/2 MS medium agar plates with 1% sucrose and imbibed at 4°C for 2 days. The plates were then placed at 21°C under a 16-h light/8-h dark photoperiod.

### Chemical treatment

Chemical treatments were performed by transferring seedlings (typically 7-day-old unless otherwise indicated) into liquid 1/2 MS medium containing 50 μM DNP (Sigma-Aldrich, Cat # D198501) or 1 μM ConA (MedChemExpress, Cat # HY-N1724). Equivalent concentrations of the solvent dimethyl sulfoxide (DMSO) were used as controls in all experiments where applicable.

In 2,4-dinitrophenol (DNP) treatment and recovery assays, Arabidopsis seedlings were grown on 1/2 MS medium agar plates for 7 days and then transferred into liquid 1/2 MS medium supplemented with either DMSO (control) or 50 μM DNP for 4 h. Following treatment, the seedlings were transferred onto fresh 1/2 MS medium agar plates and allowed to recover for 2 days. Images were captured using a digital camera, and root length was measured using ImageJ software.

### Chlorophyll extraction and measurement

Arabidopsis seedlings (fresh weight ∼ 10 mg) were collected and incubated in 1 mL of 95% ethanol for 48 h in the dark at 4°C to extract chlorophyll. The extracts were centrifuged at 12,000 rpm for 10 min at 4°C, and the absorbance of the supernatant was measured at 665 nm and 649 nm. Total chlorophyll concentration was calculated using the following equation: total chlorophyll (mg/L) = 6.63*A*_665_ + 18.08*A*_649_.

### Plasmid construction

The mitochondria-targeting sequence (Mito) derived from yeast *COXIV* transit-peptide (5’-ATGTTGTCACTACGTCAATCTATAAGATTTTTCAAGCCAGCCACAAGAACTTTGTG TAGCTCTAGATATCTGCTTGAGCAAAAACCC-3’) (Kohler *et al*., 1997) was synthesized and cloned into the *Xba*I and *Sal*I sites of pCAMBIA1300-GFP to generate pCAMBIA1300-35S::Mito-GFP vector. The coding sequences (CDS) of *ATG8a*, *ATG8e*, *GSNOR1*, *IDH1*, *VDAC1*, *VDAC2*, *VDAC3*, and *VDAC4* were PCR-amplified from wild-type Col-0 cDNA. The resulting *IDH1* and *ATG8a* CDS were cloned into pCAMBIA3301-35S-mCherry and pCAMBIA1300-35S-GFP to generate pCAMBIA3301-35S::IDH1-mCherry and pCAMBIA1300-35S::GFP-ATG8a vectors, respectively.

For recombinant protein expression, the CDS of *GSNOR1* was cloned into the *Sal*I and *Eco*RI sites of pET28a or pGEX-4T-1 to generate pET28a-GSNOR1 and pGEX-4T-1-GSNOR1 vectors, respectively. Site-directed mutagenesis was performed using appropriate PCR primer pairs to generate the following constructs: pGEX-4T-1-GSNOR1^Y152A^, pGEX-4T-1-GSNOR1^V155A^, pGEX-4T-1-GSNOR1^Y152A^ ^V155A^, pGEX-4T-1-GSNOR1^F269A^, pGEX-4T-1-GSNOR1^I272A^, and pGEX-4T-1-GSNOR1^F269A^ ^I272A^. The CDS of *VDAC1*, *VDAC2*, *VDAC3*, and *VDAC4* were cloned into the *Bam*HI and *Not*I sites of pGEX-4T-1 to generate pGEX-4T-1-VDAC1, pGEX-4T-1-VDAC2, pGEX-4T-1-VDAC3, and pGEX-4T-1-VDAC4 vectors. Truncated mutants of VDAC3 were generated using appropriate PCR primer pairs to produce pGEX-4T-1-VDAC3^D1^ through pGEX-4T-1-VDAC3^D8^ constructs. The *VDAC2* and *VDAC3* CDS were cloned into the *Bam*HI and *Sac*I sites of pMal-C2X to generate pMal-C2X-VDAC2 and pMal-C2X-VDAC3 vectors. The CDS of *ATG8a* and *ATG8e* were cloned into the *Bam*HI and *Sal*I sites of pET28a or pGEX-4T-1 to generate pET28a-ATG8a, pET28a-ATG8e, and pGEX-4T-1-ATG8a vectors. Site-directed mutagenesis was performed to generate pGEX-4T-1-ATG8a^YL65AA^ and pGEX-4T-1-ATG8a^IFV92AAA^ constructs.

For luciferase complementation assays, CDS of *GSNOR1*, *VDAC1*, *VDAC2*, *VDAC3*, and *VDAC4* were cloned into *Kpn*I and *Sal*I sites of pCAMBIA1300-NLUC or pCAMBIA1300-CLUC to generate pCAMBIA1300-GSNOR1-NLUC, pCAMBIA1300-CLUC-VDAC1, pCAMBIA1300-CLUC-VDAC2, pCAMBIA1300-CLUC-VDAC3, and pCAMBIA1300-CLUC-VDAC4 vectors.

For bimolecular fluorescence complementation (BiFC) assays, CDS of *ATG8a*, *VDAC1*, *VDAC2*, *VDAC3*, and *VDAC4* were cloned into pCAMBIA1300-35S-YCE or pCAMBIA1300-35S-YNE to generate pCAMBIA1300-35S-YCE-ATG8a, pCAMBIA1300-35S-YNE-VDAC1, pCAMBIA1300-35S-YNE-VDAC2, pCAMBIA1300-35S-YNE-VDAC3, and pCAMBIA1300-35S-YNE-VDAC4 vectors.

For subcellular localization assays in protoplast, the *TagBFP* sequence was PCR-amplified from the pLL3.7-TagBFP plasmid and cloned into pUC to generate the pUC-35S-TagBFP vector. CDS of *ATG8a*, *ATG8e*, *GSNOR1*, *IDH1*, and *VDAC3* were cloned into pUC-35S-eGFP, pUC-35S-mCherry, or pUC-35S-TagBFP to generate pUC-35S::eGFP-ATG8a, pUC-35S::eGFP-ATG8e, pUC-35S::IDH1-eGFP, pUC-35S::GSNOR1-mCherry, pUC-35S::FLAG-mCherry-VDAC3, and pUC-35S::HA-TagBFP-ATG8a vectors. Site-directed mutagenesis was performed to generate pUC-35S::GSNOR1^Y152A^ ^V155A^-mCherry and pUC-35S::GSNOR1^I272A^-mCherry constructs.

All primers used in this study are listed in Supplementary Table 2.

### Mitochondrial DNA copy number analysis

Mitochondrial DNA (mtDNA) copy number was determined by quantitative real-time PCR (qPCR) as previously described with minor modifications (Cirotti *et al*., 2021; Preuten et al., 2010). Total plant DNA was extracted using CTAB extraction buffer. Approximately 5 ng of total DNA was used as templates for each qPCR reaction. qPCR was performed using primers specific for mitochondrial genes and nuclear reference genes with UltraSYBR Mixture (CoWin Biotech, Cat # 0957) on a CFX96 Real-Time PCR Detection System (Bio-Rad). The relative mtDNA copy number was calculated by normalizing mitochondrial DNA copy number to nuclear DNA copy number (the nuclear-encoded single-copy gene *AtRpoTp*), using the 2^−ΔΔCt^ method. All primers used for qPCR are listed in Supplementary Table 2.

### Recombinant protein expression and purification

The recombinant protein expression vectors were transformed into *Escherichia coli* strain BL21 (DE3). Bacterial cultures were grown in LB medium at 37°C until OD₆₀₀ reached 0.6– 0.8, and protein expression was induced with 0.1 mM isopropyl-β-D-thiogalactoside (IPTG) at 16°C overnight. Cells were harvested by centrifugation and lysed. His-tagged, GST-tagged, and MBP-tagged recombinant proteins were purified using Ni-NTA agarose (QIAGEN, Cat # 30230), Glutathione Sepharose 4B (GE Healthcare, Cat # 17-0756-01), and Amylose Resin (New England Biolabs, Cat # E8021), respectively, according to the manufacturers’ instructions.

### Antibodies and immunoblotting

The anti-GSNOR1 antibody (also known as anti-PAR2 antibody) was described previously (Chen *et al*., 2009; Zhan *et al*., 2018). The following antibodies were obtained from commercial sources: anti-ATG8 (Agrisera, Cat # AS142811), anti-COXII (Agrisera, Cat # AS04053A), anti-GFP (Invitrogen, Cat # MA5-15256), anti-GST (CoWin Biotech, Cat # CW0084), anti-His (Engibody, Cat # AT0025), anti-HSP70 (Agrisera, Cat # AS08347), anti-HSP82 (Beijing Protein Innovation, Cat # AbM51099-31-PU), anti-MBP (EMAR, Cat # EM33019), and anti-VDAC (Agrisera, Cat # AS07212). Immunoblotting was performed as previously described (Chen *et al*., 2009; Zhan *et al*., 2018). Quantification of the immunoblot signals was performed using ImageJ software.

### GST pull-down assay

GST pull-down assays were performed as previously described (Ren et al., 2013) with minor modifications. Briefly, purified GST or GST-fusion proteins were incubated with purified His-tagged or MBP-tagged proteins and Glutathione Sepharose 4B beads for 2 h at 4°C in pull-down buffer [137 mM NaCl, 2.7 mM KCl, 10 mM Na_2_HPO_4_, 1.8 mM KH_2_PO_4_, pH 7.4, 0.05% octylphenyl-polyethylene glycol (brand name: IGEPAL CA-630)]. Beads were washed six times with pre-chilled pull-down buffer. Bound proteins were eluted by boiling in 2× SDS loading buffer and analyzed by immunoblotting.

### Co-immunoprecipitation (Co-IP) assay

Co-IP experiments were performed as previously described (Ren *et al*., 2013) with modifications. Briefly, 2-week-old seedlings were ground in liquid nitrogen and homogenized in IP buffer [50 mM Tris-HCl (pH 7.4), 1.5 mM MgCl_2_, 100 mM NaCl, 10% glycerol, 0.1% octylphenyl-polyethylene glycol (brand name: IGEPAL CA-630)] supplemented with Protease Inhibitor Cocktail (Sigma, Cat # P9599). After centrifugation at 16,000 *g* for 20 min at 4°C, supernatants were collected and incubated with anti-GFP-Nano-Agarose beads (LABLEAD, Cat # GNA-20-400) or anti-FLAG M2 Affinity Gel (Sigma-Aldrich, Cat # A2220) for 2 h at 4°C. Beads were washed five times with IP buffer, boiled in 2× SDS loading buffer, and the eluted proteins were analyzed by immunoblotting.

### Luciferase complementation imaging (LCI) and bimolecular fluorescence complementation (BiFC) assay

*Agrobacterium tumefaciens* strain GV3101 carrying the indicated expression vectors was cultured overnight in liquid Luria-Bertani medium at 28°C with shaking. *Agrobacterial* cells were collected by centrifugation at 4000 rpm for 10 min and resuspended in infiltration buffer [10 mM 2-(*N*-morpholino)ethanesulfonic acid (MES, pH 5.7), 10 mM MgCl_2_, 200 μM acetosyringone] to an OD_600_ of 0.8. The indicated *Agrobacterium* suspensions were incubated at room temperature for 3–4 h, mixed at a 1:1 ratio, and co-infiltrated into fully expanded leaves of *Nicotiana benthamiana* using a needleless syringe. For the luciferase complementation imaging (LCI) assay, *Agrobacterium*-mediated transient expression was performed as previously described (Chen *et al*., 2020). Infiltrated plants were grown under a 16-h light/8-h dark photoperiod at 21°C for 2 days. The detached leaves were sprayed with 1 mM beetle luciferin (Promega, Cat # E1605) and the luminescence signal was captured using a NightSHADE LB 985 in vivo Plant Imaging System. For the BiFC assay, the infiltrated leaves were excised 2–4 days post-infiltration and fluorescence signals were observed under a confocal laser scanning microscope (ZEISS LSM980) with excitation at 514 nm and emission at 517–595 nm for YFP.

### Yeast two-hybrid assay

The yeast two-hybrid assay was performed using the Matchmaker Gold Yeast Two-Hybrid System (Clontech Laboratories, Cat # 630466) following the manufacturer’s instructions. Briefly, the CDS of GSNOR1 were cloned into the pGBKT7 vector as the bait to screen a Mate & Plate Library-Universal Arabidopsis library (TaKaRa, Cat # 630487). The bait construct was transformed into the yeast strain Y2HGold and mated with the library strain. The mating culture was incubated at 30°C for 20–24 h with gentle shaking at 40 rpm, followed by plating on selective medium SD/-Leu/-Trp/-His/-Ade supplemented with X-α-Gal (40 mg/L) and Aureobasidin A (200 μg/L). For the yeast two-hybrid pairwise assay, the CDSs of VDAC1, VDAC2, VDAC3, and VDAC4, as well as the truncated variants of VDAC3, were cloned into the pGADT7 vector and co-transformed with pGBKT7-GSNOR1 into the Y2HGold strain using the Yeastmaker Yeast Transformation System 2 (Clontech Laboratories, Cat # 630439) according to the manufacturer’s instructions. To assess protein-protein interactions, yeast transformants were grown and serially diluted to 10^-1^, 10^-2^, and 10^-3^, and 5 μL of each dilution was spotted onto SD/-Leu/-Trp medium or SD/-Leu/-Trp/-His/-Ade medium supplemented with X-α-Gal and Aureobasidin A. Images were captured after incubation at 30°C for 3 days.

### Arabidopsis protoplast preparation and transformation

Arabidopsis protoplast preparation and transformation were performed as previously described with minor modifications (Yoo et al., 2007). Briefly, protoplasts were prepared from Arabidopsis plants grown in soil for 3–4 weeks at 21°C under a 12-h light/12-h dark photoperiod. Leaves were cut into 0.5–1-mm strips and incubated in enzyme solution [0.4 M mannitol, 20 mM MES (pH 5.7), 1.5% (w/v) cellulase R10, 0.4% (w/v) macerozyme R10, 20 mM KCl, 10 mM CaCl_2_, 0.1% (w/v) BSA] at room temperature for 3–4 h in the dark with gentle shaking at 50 rpm. The digestion mixture was diluted with an equal volume of W5 solution [154 mM NaCl, 125 mM CaCl_2_, 5 mM KCl, 2 mM MES (pH 5.7)], filtered through a 35-μm nylon mesh, and centrifuged at 100 × *g* for 2 min to collect the protoplasts. The protoplasts were washed twice with W5 solution, kept on ice for 30 min, and then resuspended in MMG solution [0.4 M mannitol, 15 mM MgCl_2_, 4 mM MES (pH 5.7)] at a concentration of 1 × 10^6^ cells/mL. For polyethylene glycol (PEG)-mediated transformation, 10 μg of the indicated plasmid DNA was mixed with 100 μL of protoplast suspension, and 110 μL of PEG solution [0.2 M mannitol, 100 mM CaCl_2_, 40% (w/v) PEG 4000] was then added. The mixture was incubated at room temperature for 10 min and then washed twice with W5 solution. The protoplasts were resuspended in 1.5 mL of W5 solution and incubated overnight at 23°C in the dark.

### Confocal microscopy and image processing

Confocal fluorescence images were acquired using an LSM980 laser scanning confocal microscope equipped with a 63× oil-immersion objective lens (Carl Zeiss). Seven-day-old Arabidopsis seedlings were treated with DMSO (control), 50 μM DNP (Sigma-Aldrich, Cat # D198501), or 1 μM ConA (MedChemExpress, Cat # HY-N1724) for the indicated times in 1/2 MS liquid medium before imaging. Mitochondria in root cells were stained with 500 nM tetramethylrhodamine ethyl ester (TMRE; Invitrogen, Cat # T669) or MitoTracker Red (MTR; Thermo Fisher, Cat # M7512) for 10 min. For visualizing depolarized mitochondria in cotyledon cells, samples were stained with TMRE for 15 min before confocal microscopy imaging. Sequential acquisition mode was applied when imaging multiple fluorescent proteins. The excitation and emission wavelengths were as follows: GFP, 488 nm excitation and 490–550 nm emission; TMRE and MitoTracker Red, 561 nm excitation and 580–630 nm emission; mCherry, 561 nm excitation and 580–630 nm emission; TagBFP, 405 nm excitation and 410–585 nm emission; YFP, 514 nm excitation and 517–595 nm emission. Images were processed and analyzed using ZEN software.

### Transmission electron microscopy (TEM)

Samples were immediately fixed in 2.5% (v/v) glutaraldehyde in 0.1 M phosphate buffer (pH 7.2) for 2 h at room temperature, and then rinsed three times with 0.1 M phosphate buffer (pH 7.2) for 20 min each. Samples were post-fixed in 1% (v/v) osmium tetroxide (OsO_4_) for 2 h, followed by three rinses with phosphate buffer (15 min each). The samples were then dehydrated through a graded acetone series [30%, 50%, 70%, 90%, and 100% (three times); v/v in ddH_2_O] at room temperature, with 20 min incubation at each concentration. Dehydrated samples were infiltrated with acetone/EMBed-812 resin (Electron Microscopy Sciences, Cat #14120) mixture at ratios of 3:1, 1:1, 1:3 (v/v) for 12 h each, followed by 100% EMBed-812 resin for 12 h. Samples were embedded in fresh EMBed-812 resin and polymerized at 37°C for 12 h, 45°C for 12 h, and 60°C for 48 h. Ultrathin sections (70 nm) were prepared using an ultramicrotome (Leica, EM UC7). Sections were observed and photographed using a transmission electron microscope (Hitachi, HT-7700) at an accelerating voltage of 80 kV.

### Quantification and statistical analysis

For quantification analysis, the mean SD or SEM were calculated and compared to the control. The significance (*P* value) was determined using two-tailed Student’s *t*-test or One-way ANOVA test (specified in Figure legends). \**P* < 0.05, \*\**P* < 0.01, \*\*\**P* < 0.001. All experiments were repeated at least 3 times, and representative results are shown.

## Data availability

All other data are available in the main text or supplementary materials.

## Acknowledgements

We thank Dingzhong Tang, Yikun He, and Ye Tian for kindly providing seeds and plasmids; Yanbao Tian for help on confocal microscopy and transmission electron microscopy; Byung-Ho Kang for sharing unpublished data; members of Zuo Lab for discussion and suggestions. This work was supported by grants from the National Natural Science Foundation of China (NSFC; 31830017 and 32200256) and State Key Laboratory of Plant Genomics (SKLPG2023-22).

## Author Contributions

H.L. and H.G. performed most of the experiments, assisted by R.W., W.W., P.-F.J., S.S., and R.B. J.Z., W.-C.Y., and H.-J.L. supervised the research. J.Z., H.L., H.G., W.-C.Y., and H.-J.L. analyzed the data. J.Z. and H.L. wrote the manuscript, assisted by H.G. All authors read and approved the manuscript.

## Ethics declarations Competing interests

The authors declare no competing interests.

## Supplementary information

**Supplementary Table 1.** Putative GSNOR1-interacting proteins identified by yeast two-hybrid screen.

**Supplementary Table 2.** Primers used in this study.

**Supplementary Video 1.** Co-localization of GSNOR1-GFP and IDH1-mCherry.

## Extended data

**Extended Data Fig. 1:**
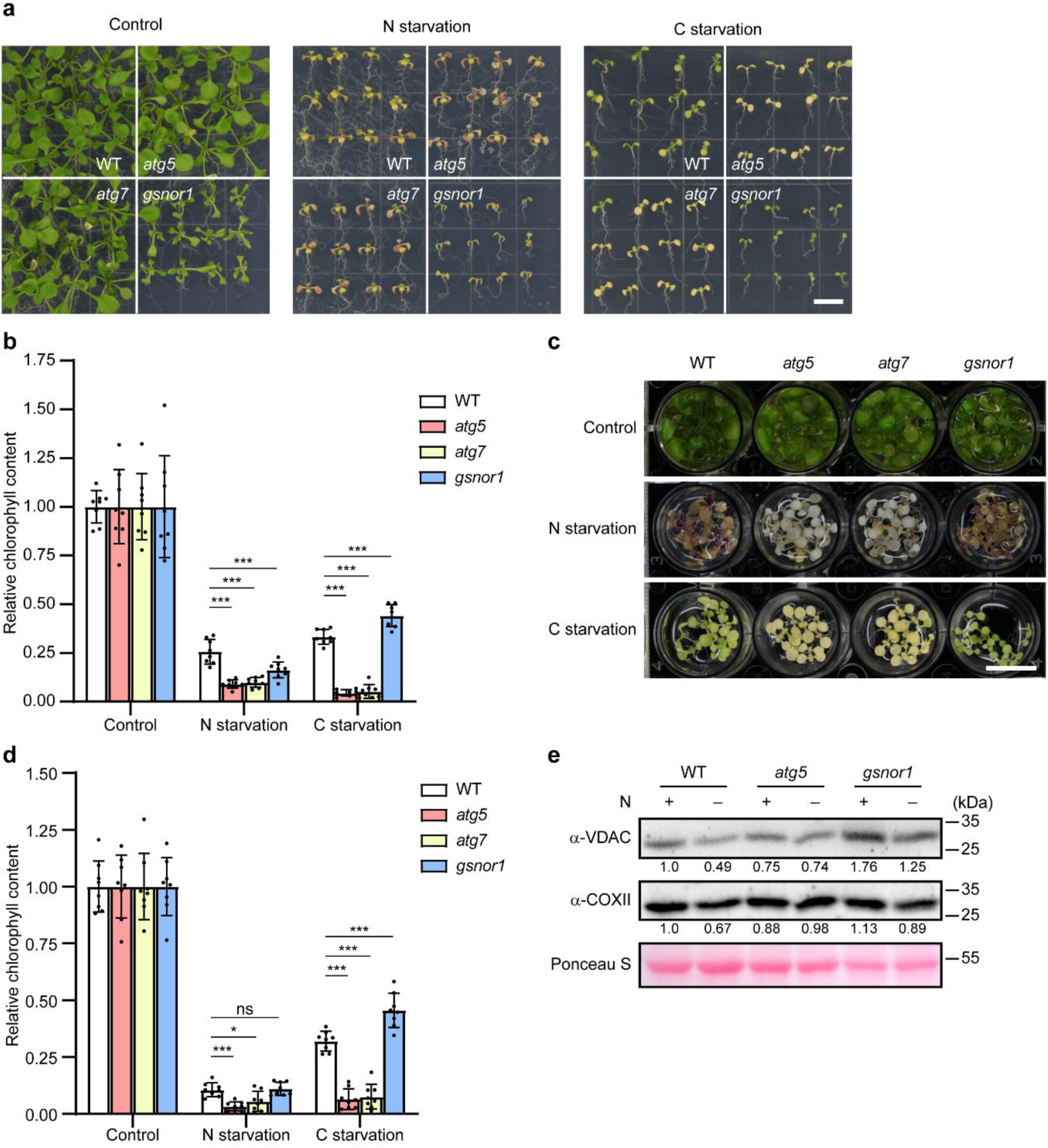
GSNOR1 is not involved in starvation-induced mitochondrial degradation. **a,** Seven-day-old wild-type (WT; Col-0), *atg5*, *atg7*, and *gsnor1* seedlings germinated and grown on 1/2 MS medium were transferred to fresh 1/2 MS medium (control), nitrogen-free medium (N starvation), or sucrose-free agar medium cultured in the dark (C starvation). Seedlings were cultured for additional 8 days prior to photographing. Scale bar, 1 cm. **b,** Analysis of chlorophyll contents of seedlings shown in (**a**). The relative chlorophyll content of each genotype under control conditions was set as 1.0. Data are presented as mean ± SD of eight biological replicates. Statistical significance was determined by one-way ANOVA followed by Tukey’s multiple comparisons test. \*\*\**P* < 0.001. **c,** Seven-day-old wild type (WT; Col-0), *atg5*, *atg7*, and *gsnor1* seedlings germinated and grown on 1/2 MS medium were transferred to fresh liquid 1/2 MS medium (control), nitrogen-free medium (N starvation), or sucrose-free medium cultured in the dark (C starvation). Seedlings were cultured for additional 8 days prior to photographing. Scale bar, 1 cm. **d,** Analysis of chlorophyll contents of seedlings shown in (**c**). The relative chlorophyll content of each genotype under control conditions was set as 1.0. Data are presented as mean ± SD of eight biological replicates. Statistical significance was determined by one-way ANOVA followed by Tukey’s multiple comparisons test. ns, not significant; \**P* < 0.05; \*\*\**P* < 0.001. **e,** Analysis of mitochondrial outer membrane proteins VDACs and the mitochondrial inner membrane protein COXII in wild-type (WT), *atg5*, and *gsnor1* seedlings by immunoblotting. Seven-day-old seedlings germinated and grown on 1/2 MS agar medium were transferred to liquid 1/2 MS medium or nitrogen-free medium for 24 h. Equal loading was shown by Ponceau S staining. Quantification of band intensities is shown below the blot. The relative protein level in WT grown under control conditions was set as 1.0.

**Extended Data Fig. 2:**
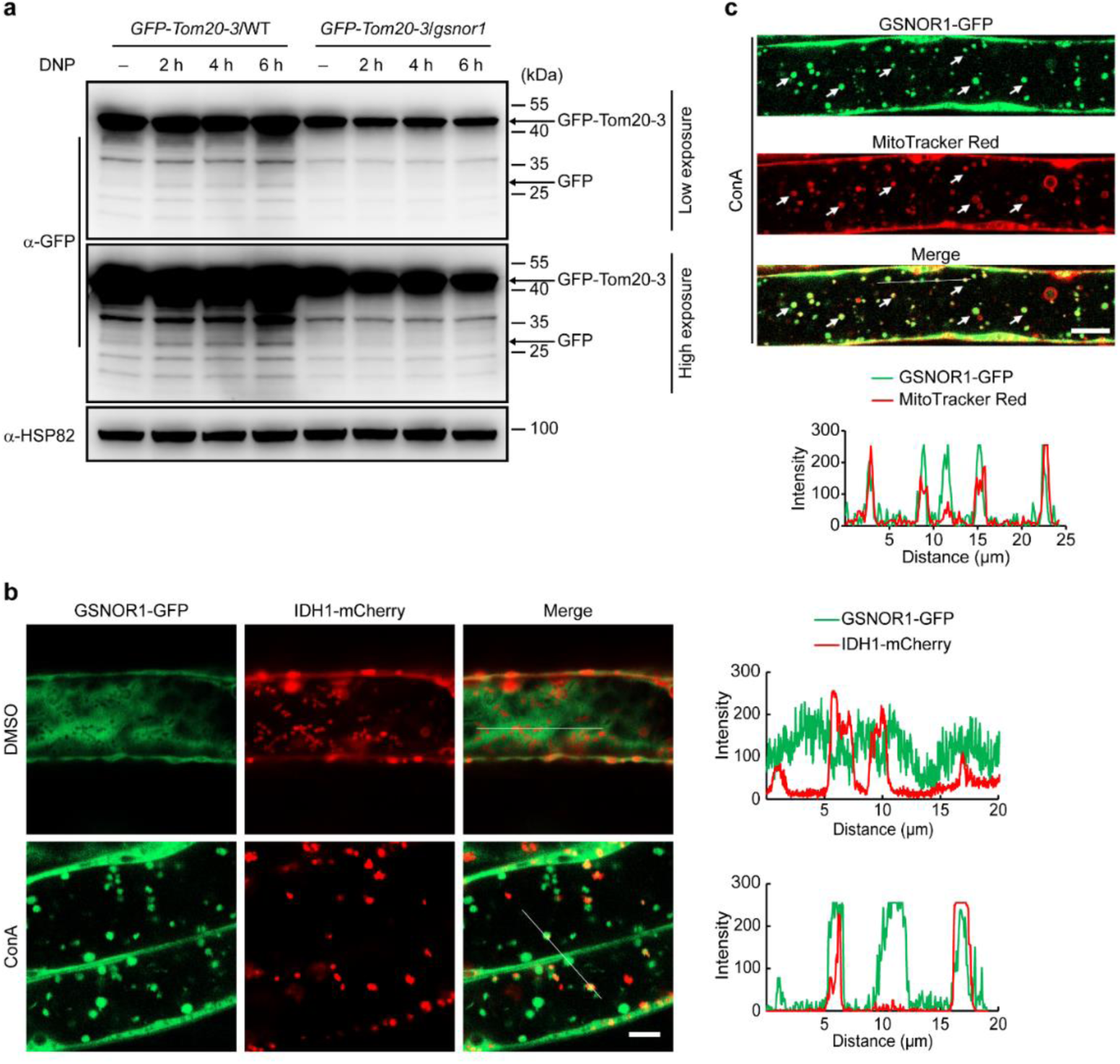
GSNOR1 localizes to depolarized mitochondria. **a,** Analysis of mitophagic flux in *GFP-TOM20-3* transgenic seedlings in the wild-type (WT) and *gsnor1* genetic backgrounds. Cleavage of GFP from GFP-TOM20-3 was detected using an anti-GFP antibody. Arrows indicate the bands of full-length GFP-TOM20-3 and free GFP, respectively. Seven-day-old transgenic seedlings were treated with DMSO (control, 2 h) or with 50 µM DNP for the indicated times. HSP82 served as a loading control. **b,** Confocal microscopy of roots from 7-day-old seedlings carrying both *GSNOR1-GFP* and *IDH1-mCherry* transgenes. The samples were treated with DMSO or ConA (1 µM) for 12 h prior to imaging. Line scans on the right side of the images indicate co-localization between GSNOR1-GFP (green) and IDH1-mCherry (red). Scale bar, 5 µm. **c,** Confocal microscopy of roots from 5-day-old *GSNOR1-GFP* transgenic seedlings treated with 1 µM ConA for 12 h, followed by staining with MitoTracker Red for 10 min. Line scans below the images indicate co-localization between GSNOR1-GFP (green) and MitoTracker Red (red). White arrows indicate punctate co-localization signals. Scale bar, 10 μm.

**Extended Data Fig. 3:**
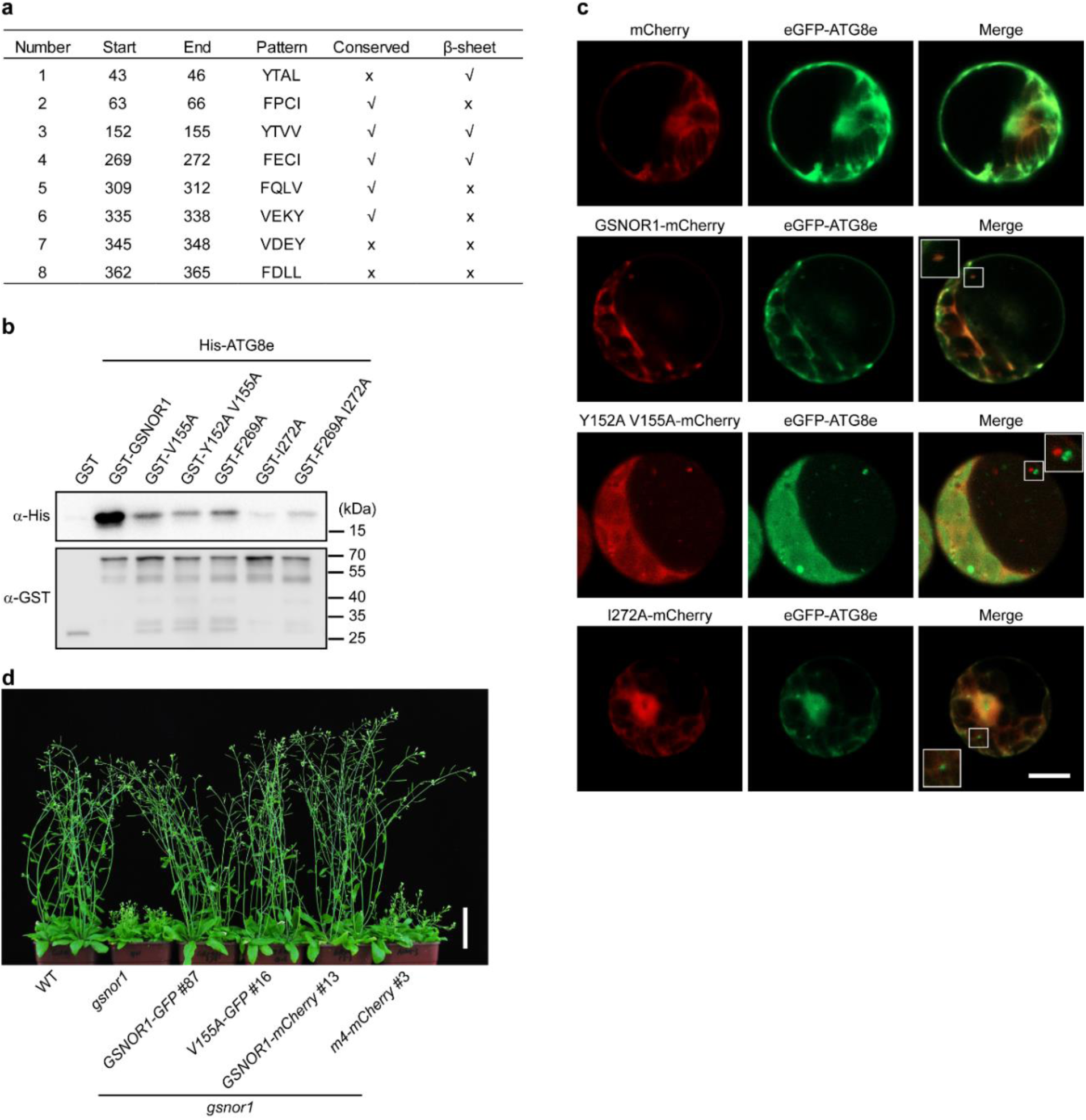
GSNOR1 interacts with ATG8 via its AIM motifs. **a,** Putative AIM motifs of GSNOR1. The search was performed using the hfAIM tool as described(Xie et al., 2016). **b,** Analysis of the interaction of His-ATG8e and GST-GSNOR1 recombinant proteins by a GST pull-down assay. GST protein was used as a negative control. **c,** Confocal microscopy of Arabidopsis protoplasts transformed with *GSNOR1-mCherry* and *eGFP-ATG8e* and then cultured for 24 h prior to imaging. Scale bar, 10 µm. **d,** Six-week-old plants with the indicated genotypes grown under long-day condition (16-h light/8-h dark) in a growth chamber. See also Fig. 3e. Same plants of the control group, i.e., wild type (WT), *gsnor1*, *GSNOR1-GFP* (#87), and *GSNOR1-mCherry* (#13), were used for photographing in Fig. 3e and here. Scale bar, 5 cm.

**Extended Data Fig. 4:**
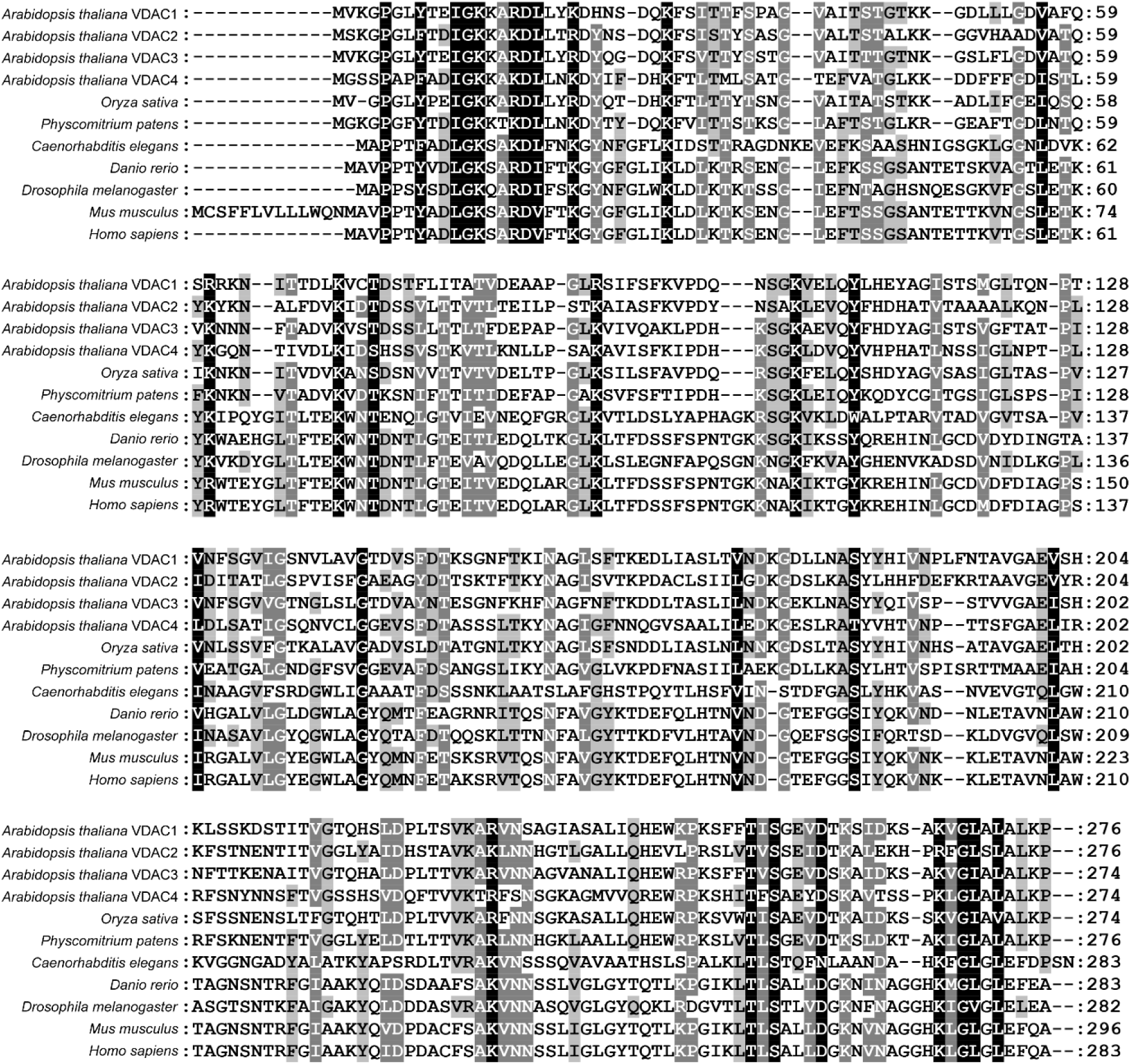
Alignment of VDAC proteins. Protein sequence alignment of VDAC family proteins. Protein sequences used for alignment with the following accession numbers: *Arabidopsis thaliana* VDAC1: NP_186777; *Arabidopsis thaliana* VDAC2: NP_201551; *Arabidopsis thaliana* VDAC3: NP_001190314; *Arabidopsis thaliana* VDAC4: NP_200557; *Oryza sativa*: NP_001390674; *Physcomitrium patens*: XP_024397563; *Caenorhabditis elegans*, NP_501211; *Danio rerio*: NP_001001404; *Drosophila melanogaster*: NP_001033899; *Mus musculus*: NP_001387916; *Homo sapiens*: NP_001387937. Sequence alignment was performed using Clustal X2, and the results were visualized in Genedoc.

**Extended Data Fig. 5:**
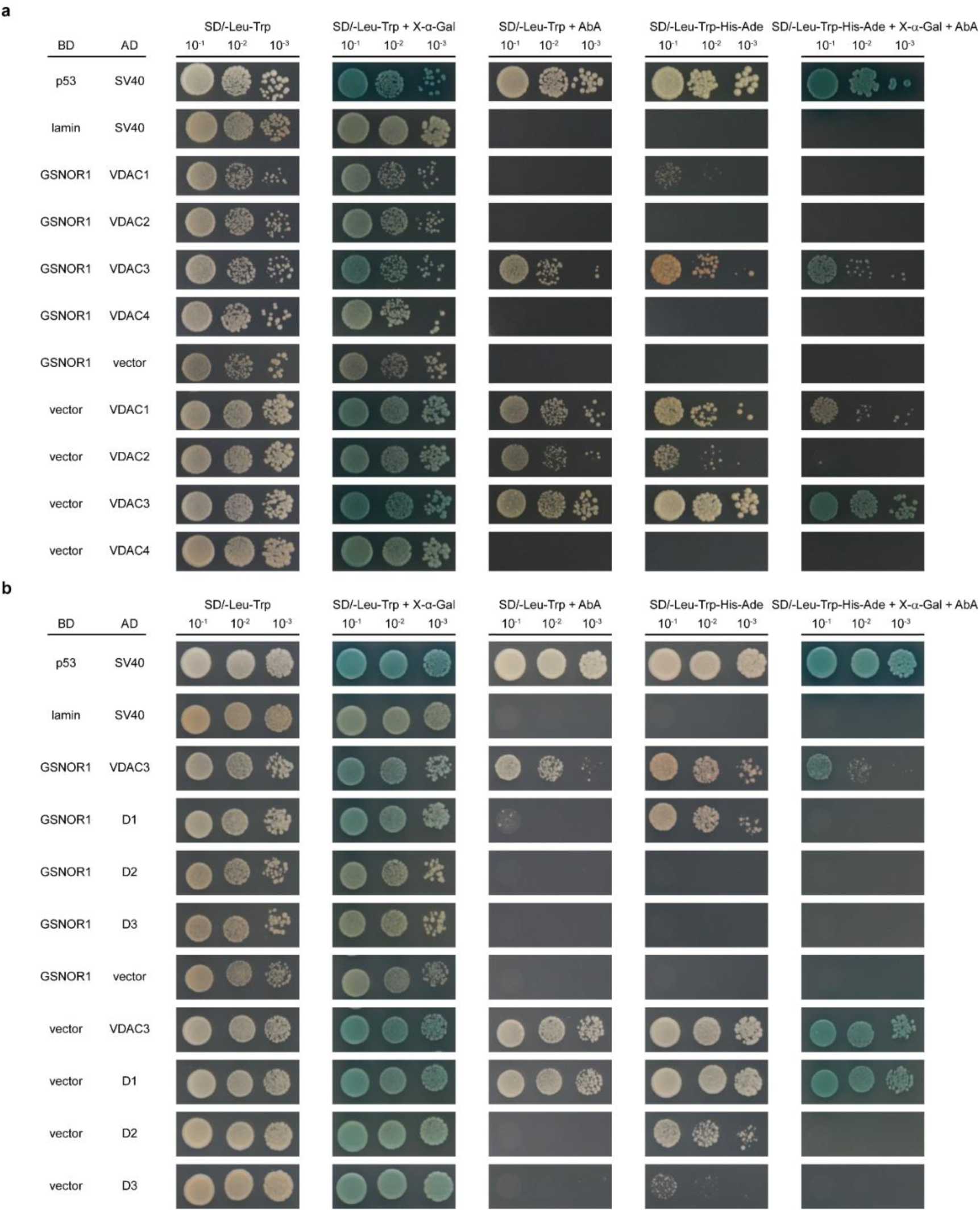
Analysis of interactions between GSNOR1 and VDAC proteins. **a,b,** Analysis of the interaction of GSNOR1 with VDAC1 through VDAC4 proteins (**a**) or truncated VDAC3 mutant proteins (**b**) by the yeast two-hybrid assay. GSNOR1 is fused to the GAL4 DNA-binding domain (BD), and VDAC proteins are fused to the GAL4 activation domain (AD). The paired combinations as indicated are transformed and expressed in yeast strain Y2HGold. Protein interactions were assessed by activation of the reporter genes *HIS3*, *ADE2*, *AUR1-C*, or *MEL1*. The p53-SV40 interaction and lamin-SV40 interaction served as positive and negative controls, respectively. pGBKT7 and pGADT7 were used as the bait and prey vectors, respectively.

**Extended Data Fig. 6:**
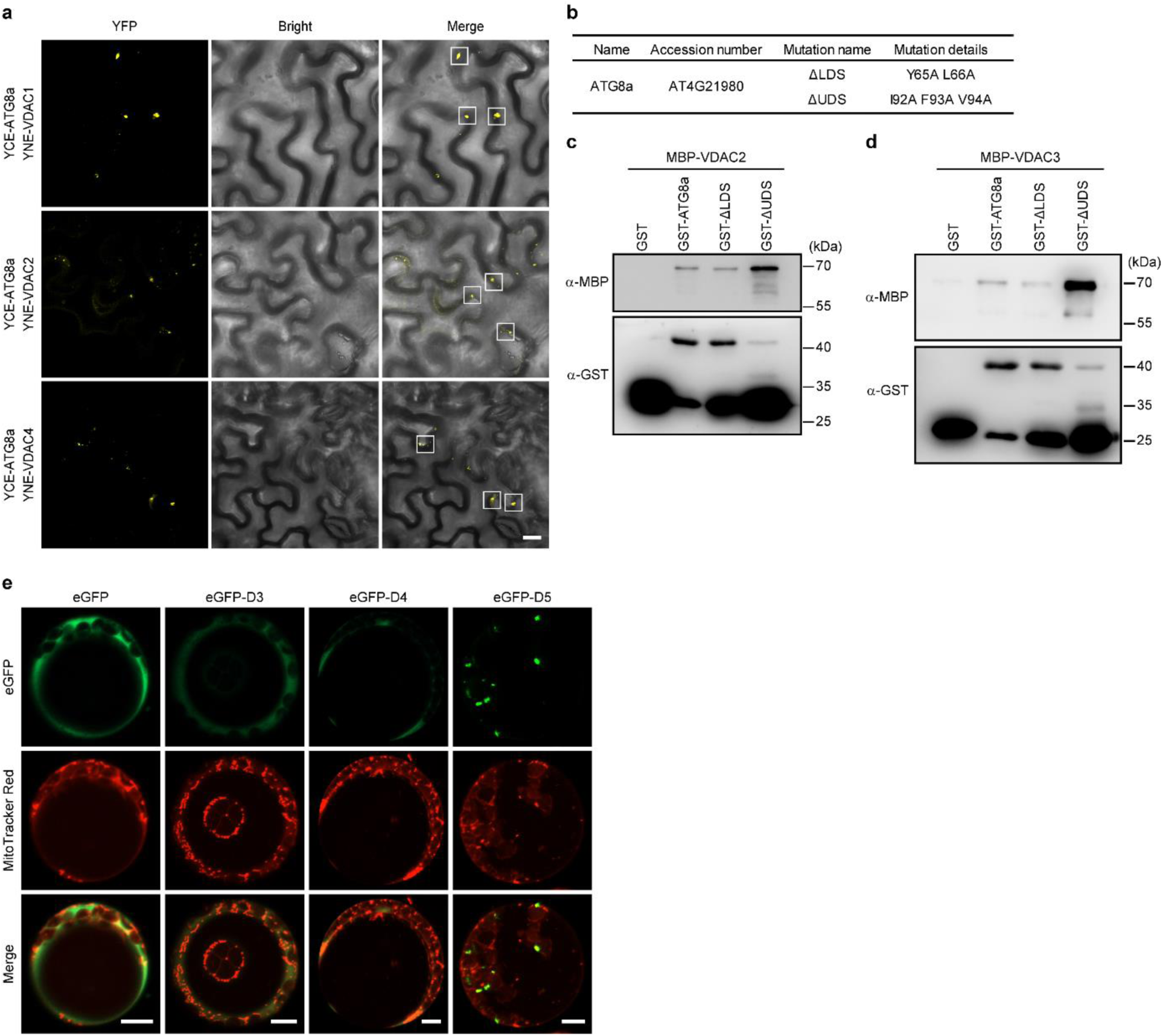
Functional characterization of VDAC proteins. **a,** Analysis of the interaction of ATG8a and VDAC proteins by a bimolecular fluorescence complementation assay in tobacco leaves infiltrated with *Agrobacterium* transformed with various combinations of transgenes as indicated. Confocal microscopy was performed 4 days post-infiltration. Co-localization signals are shown in the boxed regions. Scale bar, 20 µm. **b,** Positions and nature of mutations in UDS (UIM-docking site) and LDS (LIR/AIM-docking site) motifs of ATG8a. **c,d,** Analysis of the interaction of GST-ATG8a and MBP-VDAC2/VDAC3 recombinant proteins shown in (**b**) by pull-down assay. **e,** Confocal microscopy of Arabidopsis protoplasts transiently expressing eGFP, eGFP-D3, eGFP-D4, or eGFP-D5. Protoplasts were stained with 200 nM MitoTracker Red for 15 min prior to imaging. Scale bars, 10 µm.

